# Therapeutic Reversal of Chemically Induced Parkinson Disease by Converting Astrocytes into Nigral Neurons

**DOI:** 10.1101/2020.04.06.028084

**Authors:** Hao Qian, Jing Hu, Dongyang Zhang, Fan Meng, Xuan Zhang, Yuanchao Xue, Neal K. Devaraj, Steven F. Dowdy, William C. Mobley, Don W. Cleveland, Xiang-Dong Fu

**Author notes:** Corresponding author: Xiang-Dong Fu.

## Abstract

Parkinson disease is characterized by loss of dopamine neurons in the substantia nigra. As with other neurodegenerative diseases, no disease-modifying treatments exist. While most treatment objectives aim to prevent neuronal loss or protect vulnerable neuronal circuits, an important alternative is to replace lost neurons to reconstruct disrupted circuits. Herein we report an efficient single-step conversion of isolated mouse and human astrocytes into functional neurons by depleting the RNA binding protein PTB. Applying this approach to mice with a chemically induced Parkinson’s phenotype, we provide evidence that disease manifestations can be potently reversed through converting astrocytes into new substantia nigral neurons, effectively restoring dopamine levels via reestablishing the nigrostriatal dopamine pathway. We further demonstrate similar disease reversal with a therapeutically feasible approach using antisense oligonucleotides to transiently suppress PTB. These findings identify a generalizable therapeutic strategy for treating neurodegenerative disorders through replacing lost neurons in the brain.

## INTRODUCTION

Regenerative medicine holds great promise for addressing disorders that feature cell loss. While cell replacement has enjoyed remarkable success in treating hematopoietic disorders^1^, in other diseases this approach has shown either limited efficacy or association with risk of triggering immune responses and/or tumor formation. Given increasing evidence for cellular plasticity of differentiated somatic cells^2^, *trans*-differentiation approaches for *in situ* switching of cell fate have gained significant momentum^3^. Particularly noteworthy has been the successful conversion of fibroblasts to cardiomyocytes in the heart^4,5^ and non-neuronal cells to neurons in the brain^6,7^.

While progress in regenerative medicine is encouraging, few studies have achieved its ultimate goal: disease reversal. Heinrich and colleagues^8^ outlined five major milestones for *in vivo* reprogramming: (i) identifying source cell types, (ii) developing the best strategy for conversion, (iii) matching the phenotype of a desirable cell type, (iv) overcoming environmental constraints in the host, and most importantly, (v) promoting functional integration to restore lost function with measurable benefits at the organismal level. In Parkinson disease (PD) models, for example, employing methods to convert endogenous non-neuronal cells into neurons has yet to fully achieve these milestones.

In the mouse brain, non-neuronal cell types, including oligodendrocytes and astrocytes, show sufficient plasticity for reprogramming^6,9,10^. Studies have also provided evidence for functional integration of converted neurons^6,7,10^, including behavioral benefits after converting striatal astrocytes into dopaminergic neurons^7^. However, since dopaminergic neurons originate from substantia nigra, it has yet to be shown that *in vivo* converted neurons reconstitute the nigrostriatal dopamine pathway, a milestone probably critical for long-term functional recovery in PD.

The vast majority of current *in vivo* reprogramming approaches rely on one or more lineage-specific transcription factors (TFs)^6,7,10^, whose functional impact in most cases was empirically defined. In contrast to TF-based approaches, we recently elucidated roles for the RNA binding protein PTB and its neuronal analog nPTB in two consecutive regulatory loops that control neuronal induction and maturation, and demonstrated efficient conversion of both mouse and human fibroblasts to functional neurons through sequential depletion of PTB and nPTB^11^. Importantly, sequential down-regulation of PTB and nPTB occurs naturally during neurogenesis, and once triggered, both loops are self-enforced. Rather than relying on overexpression of an exogenous gene(s), such as TFs, this strategy takes full advantage of the endogenous neurogenesis program to convert non-neurons into neurons. This on-off “switch” for neurogenesis thus appears ideal for *in vivo* reprogramming, potentiating a transient intervention to achieve long-term benefits.

Exploring this strategy for direct reprogramming of astrocytes into neurons in the mouse brain, we now report potent conversion of astrocytes into dopamine neurons in the substantia nigra, a substantial fraction of which send axonal projections into the striatum. In a chemically-induced mouse PD model, we show that this strategy effectively restores dopamine production and reverses the PD phenotype, thus satisfying all five of the Heinrich et al^8^ milestones for *in vivo* reprogramming. Furthermore, employing a chemogenetic approach, we provide evidence that converted neurons are directly responsible for the observed phenotypic recovery. Finally, we demonstrate that an antisense oligonucleotide (ASO) against PTB is effective in reversing the PD phenotype, supporting the feasibility of a transient, single-step strategy for treating PD and perhaps other neurodegenerative diseases.

## RESULTS

### Key neurogenesis factors are active in astrocytes

Astrocytes offer several advantages for *in vivo* reprogramming in the nervous system. These non-neuronal cells are abundant in brain and spinal cord^12,13^, become proliferative in response to injury^14^, and demonstrate a significant degree of plasticity in switching cell fate^15^. As we established for fibroblasts^11,16^, PTB suppresses a neuronal induction loop in which the microRNA miR-124 inhibits the transcriptional repressor REST responsible for suppressing a large number of neuronal-specific genes that include miR-124 (Fig. 1a, loop I). PTB down-regulation also induces the expression of nPTB, which is part of a second loop for neuronal maturation that includes the transcription activator Brn2 and miR-9 (Fig. 1a, loop II). Sequential down-regulation of PTB and nPTB activate both loops to generate functional neurons from fibroblasts^11^.

**Fig. 1.**
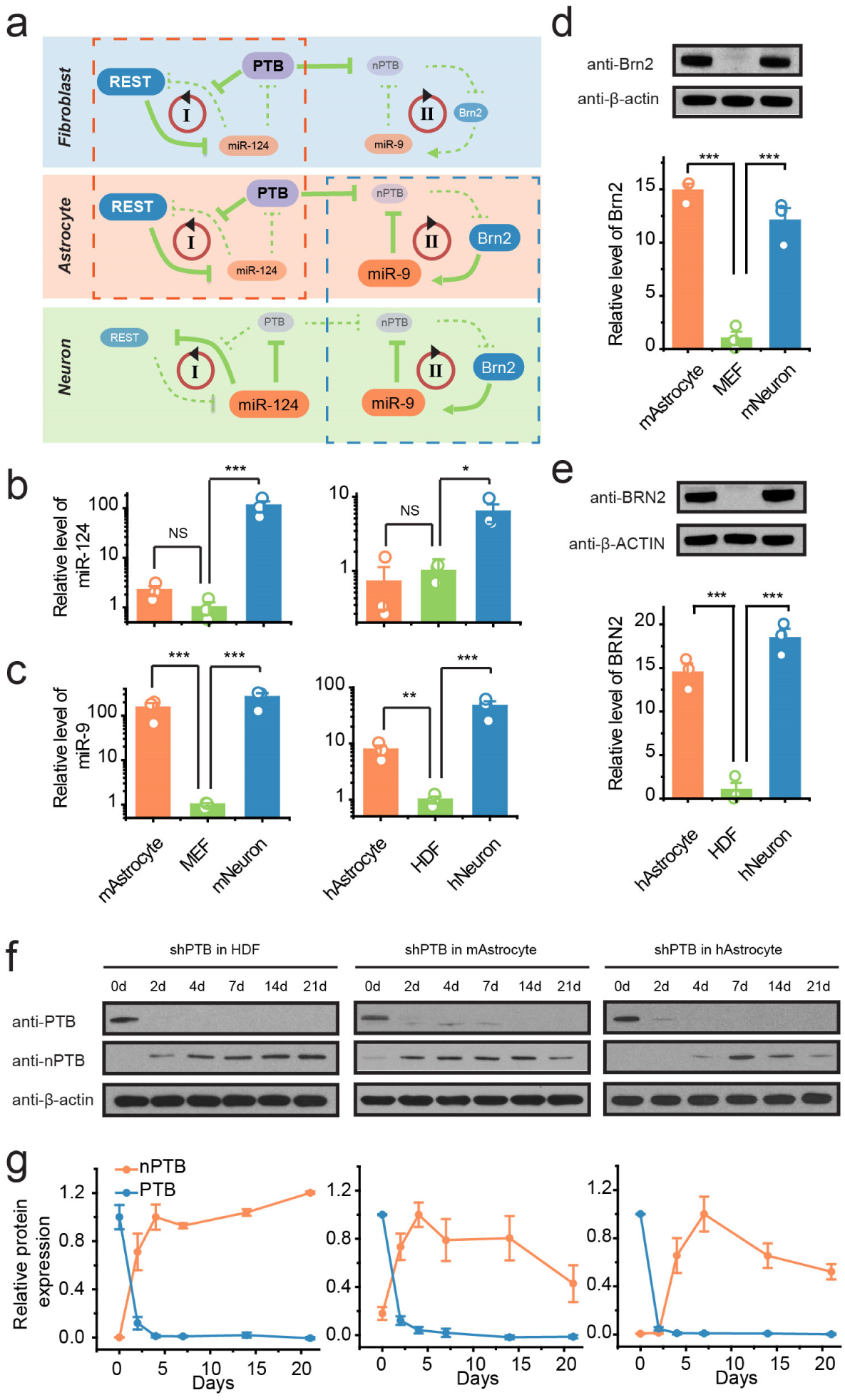
PTB down-regulation activates key factors in both neuronal induction and maturation programs in mouse and human astrocytes. **a**, Two consecutive regulatory loops controlled by PTB for neuronal induction and nPTB for neuronal maturation in fibroblast, astrocyte, and neurons. The expression levels of individual components are represented by large (for high) and small (for low) font sizes. Red dashed box highlights similarity of the first loop between fibroblast and astrocyte and blue dashed box emphasizes the similarity of the second loop between astrocyte and neuron. **b** and **c**, Expression levels of miR-124 (**b**) and miR-9 (**c**) determined by RT-qPCR in astrocytes from mouse (mAstrocyte) or human (hAstrocyte), mouse embryonic fibroblasts (MEF) and human adult fibroblasts (HDF) and neurons from embryonic mouse brain (mNeuron) or differentiated from human neural progenitor cells (hNeuron). **d** and **e**, The expression levels of Brn2 examined by immunoblotting in mouse and human astrocytes, MEF and HDF, and neurons from mouse and human. **f** and **g**, Time-course analysis of nPTB levels by immunoblotting (**f**) in response to PTB knockdown in HDF (left), mouse (middle) and human astrocytes (right). The data are quantified in **g**. Statistical results in **b-e** are represented as mean+/- SEM; *p<0.05; **p<0.01; ***p<0.001 based on ANOVA with post-hoc Tukey test (n=3 biological replicates).

To explore this RNA binding protein-regulated cascade as a strategy to convert astrocytes into neurons, we first characterized the expression of key microRNAs in primary mouse and human astrocytes. As in fibroblasts, miR-124 was expressed at a low level in both human and mouse astrocytes (Fig. 1b), which explains the high REST levels in this non-neuronal cell type^17^. Unexpectedly, as in neurons but not fibroblasts, miR-9 was highly expressed in both mouse and human astrocytes (Fig. 1c). Brn2 followed the same expression pattern as miR-9, high in both astrocytes and neurons, but low in fibroblasts (Fig. 1d, e), which is consistent with astrocytes and neurons sharing common progenitors^18^. These data demonstrate that while astrocytes resemble fibroblasts in the PTB-regulated loop (Fig. 1a, red dashed box), they are already neuron-like in the nPTB-regulated loop (Fig. 1a, blue dashed boxes).

Given that astrocytes already express miR-9 and Brn2, two key factors required for neuronal maturation, we tested the possibility that PTB knockdown-induced expression of nPTB would be immediately counteracted by miR-9, thus resulting only in transient nPTB expression, as seen during neurogenesis from neural stem cells^19^. Indeed, in contrast to human fibroblasts, PTB knockdown in mouse and human astrocytes led to transient nPTB induction (Fig. 1f, g). We reasoned, therefore, that depleting PTB alone from astrocytes already expressing high levels of endogenous Brn2 and miR-9 might potentiate their reprogramming into mature neurons.

### PTB down-regulation efficiently converts astrocytes to functional neurons *in vitro*

To test if PTB down-regulation efficiently converts astrocytes to mature neurons, we dissociated mouse astrocytes from cerebral cortex of postnatal day 4 to 5 (P4-5) pups^6^ or obtained human fetal astrocytes from a commercial source (ScienCell). Cells from both sources expressed the expected astrocytic markers GFAP and ALDH1L1, without detectable contamination of neuronal cells, as indicated by the absence of cells expressing proteins unique to neurons or neural crest progenitors (Extended Data Fig. 1).

We infected mouse astrocytes with a lentiviral vector expressing a small hairpin RNA against PTB (shPTB). Four weeks after transduction, ∼50% of shPTB-treated cells maintained in standard neuronal differentiation medium showed neuronal morphology and positive staining for the pan-neuronal markers Tuj1 and MAP2, while cells transduced with a control shRNA did not (Fig. 2a). The shPTB-induced neurons also expressed markers of mature neurons, including NeuN and a Neuron Specific Enolase (NSE) (Fig. 2b). To define the types of converted neurons, we examined markers for glutamatergic neurons (VGlut1), GABAergic neurons (GAD67), dopaminergic neurons (tyrosine hydroxylase, TH) (Fig. 2b). The majority of induced neurons were glutamatergic or GABAergic, with only a few (∼2%) Tuj1^+^ cells expressing TH (Fig. 2c). We examined the expression of additional dopaminergic markers, such as SLC6A3 and FoxA2 by RT-qPCR, as well as DAT and VMAT2 by immunostaining, and again observed their induction at low efficiency (Extended Data Fig. 2a-d). None of the induced neurons expressed detectable cholinergic or serotonergic markers, such as choline acetyltransferase or tryptophan hydroxylase (data not shown).

**Fig. 2.**
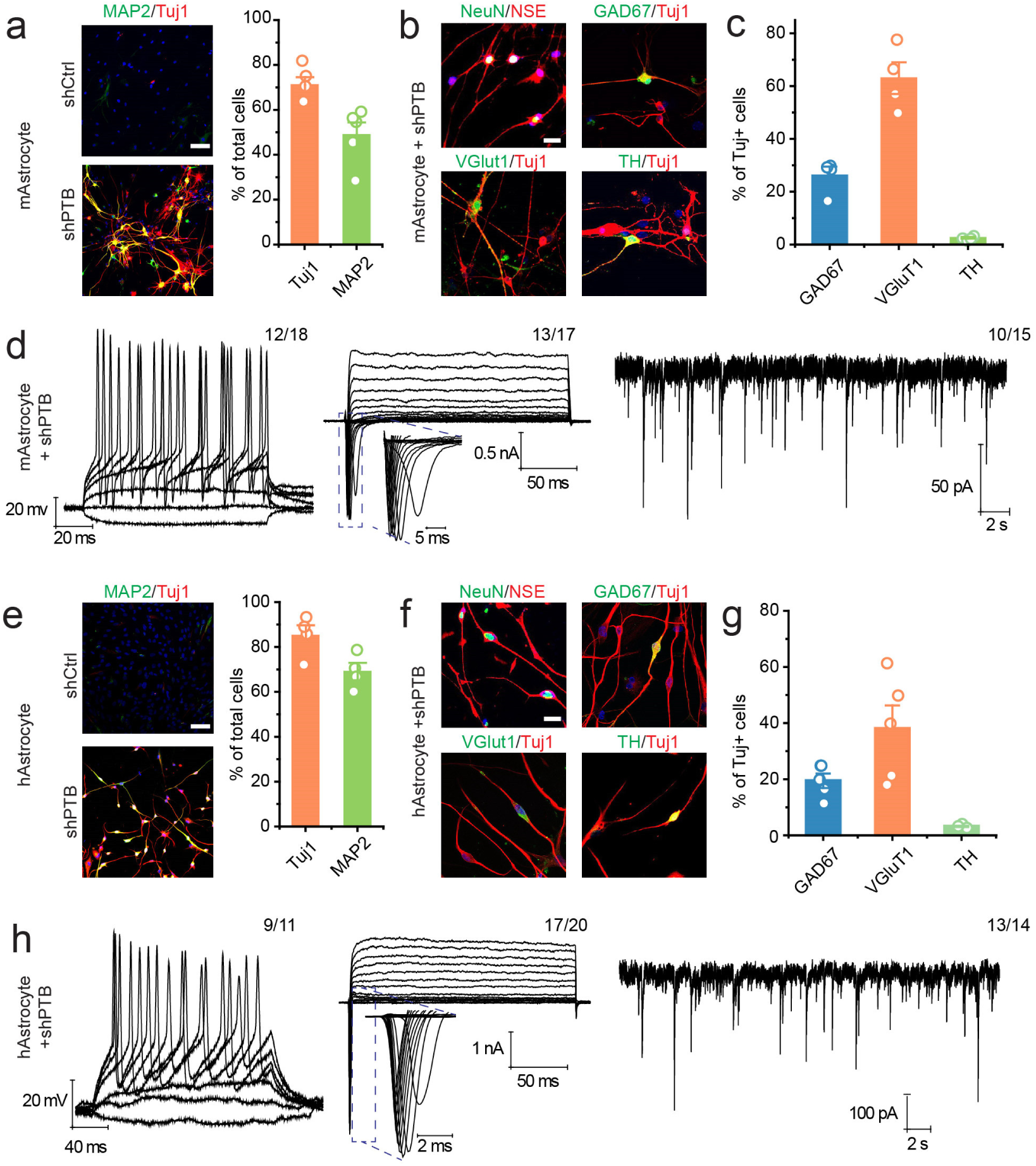
Efficient generation of functional neurons from mouse and human astrocytes by PTB knockdown. **a**, PTB knockdown-induced neurons from mouse astrocytes stained for the pan-neuronal markers Tuj1 (red) and MAP2 (green). Mouse astrocytes infected with control virus (shCtrl) showed no positive staining of neuronal markers under same culture conditions. Right, quantification based on 5 biological repeats. Scale bar: 100 μm. **b**, Characterization of induced neurons by examining the expression of markers for mature neurons (NeuN, NSE) and markers for different neuronal subtypes, glutamatergic neurons (VGlut1), GABAergic neurons (GAD67) and dopaminergic neurons (TH). Scale bar: 30 μm. **c**, Quantification for subtypes of converted neurons from mouse astrocytes. Data are based on 4 biological repeats. **d**, Electrophysiological analysis of induced neurons from mouse astrocytes, showing repetitive action potentials (left), large currents of voltage-dependent sodium and potassium channels (middle), and after co-culture with rat astrocytes, spontaneous post-synaptic currents were also recorded (right). The number of cells that showed the recorded activity versus the total number of cells examined is indicated on top right in each panel. **e**, Induced expression of Tuj1 (red) and MAP2 (green) by PTB knockdown in human astrocytes. Scale bar: 100 μm. Right panel shows the quantified data from 4 biological repeats. **f**, Induced neurons from human astrocytes expressed markers of mature neurons (NeuN, NSE) and markers of different neural subtypes (VGlut1, GAD67 and TH). Scale bar: 40um. **g**, Quantification for subtypes of converted neurons from human astrocytes. Data are based on 5 biological repeats. **h**, Electrophysiological analysis of induced neurons of induced neurons from human astrocytes, showing repetitive action potentials (left), large currents of voltage dependent sodium and potassium channels (middle) and spontaneous post-synaptic currents (right). The number of cells that showed the recorded activity versus the total number of cells examined is indicated on top right in each panel. Note that the quantitative data shown in **a, c, e, g** are intended to present the actual levels, not for emphasizing statistical differences between different groups.

Electrical activity of PTB knockdown-converted cells was measured by patch clamp recordings on individual neurons 6∼8 weeks after shPTB treatment. Most patched cells showed currents of voltage-gated sodium/potassium channels and repetitive action potential firing (Fig. 2d). When converted neurons were co-cultured with freshly isolated GFP-marked rat astrocytes, spontaneous postsynaptic events of varying frequencies were detected (Fig. 2d). These neuronal activities likely reflected the responses to synaptic inputs from both glutamatergic and GABAergic neurons because addition of antagonists of glutamatergic channel receptors [2,3-dihydroxy-6-nitro-7-sulfamoyl-benzo[f]quinoxaline-2,3-dione (NBQX) plus D(-)-2-amino-5-phosphonovaleric acid (APV)] and an antagonist of GABA_A_ channel receptors [Picrotoxin (PiTX)] sequentially blocked the signals (Extended Data Fig. 3a). No neuronal electrophysiological properties were detected in patch clamp recording of cells transduced with the control shRNA (Extended Data Fig. 3b).

Human astrocytes were also efficiently reprogrammed to neurons. Four weeks after transduction with the shPTB lentiviral vector, nearly quantitative astrocyte-to-neuron conversion was observed, with ∼90% of the cells marked by Tuj1 (Fig. 2e). Converted neurons also expressed NeuN and NSE (Fig. 2f), and, as with mouse astrocytes, human astrocytes were largely converted to glutamatergic or GABAergic neurons, with only a small percent (1 to 2%) expressing a detectable level of TH (Fig. 2f, g). Compared with mouse astrocytes, the conversion efficiency was higher with human astrocytes (compare Fig. 2a and 2e), while the relative percentages of GAD67 and VGlut1 neuron subtypes were lower (compare Fig. 2c and 2g), perhaps indicating higher diversification of human astrocyte-derived neurons. Electrophysiological measurements were performed to show the presence of currents carried by voltage-gated channels and repetitive action potentials in the vast majority of neurons within 5∼6 weeks after depletion of PTB. When co-cultured with rat astrocytes, most human astrocyte-converted neurons also exhibited spontaneous postsynaptic events (Fig. 2h), which could be sequentially blocked NBQX+APV and PiTX (Extended Data Fig. 3c). No neuronal electrophysiological properties were detected in patch clamp recording of cells transduced with the control shRNA (Extended Data Fig. 3d). Collectively, these data demonstrate that both mouse and human astrocytes can be efficiently converted to functional neurons in a single step through down-regulating PTB.

### Direct conversion of astrocytes into neurons in mouse midbrain

Recognizing the potential benefit of direct astrocyte-to-neuron conversion for restoring neurons and initiating reconstruction of neuronal circuits in the context of neurodegenerative disorders, we attempted to reprogram astrocytes into neurons directly in the brain. For this purpose, we focused on the substantia nigra of midbrain, the locus of dopaminergic neurons, whose loss is causative of PD. We designed an AAV-based strategy by using AAV serotype 2 to express shPTB for *in vivo* delivery (Fig. 3a). To enable lineage tracing of converted neurons, we placed the red fluorescence protein (RFP) gene 5’ to shPTB. To allow for conditional expression of both RFP and shPTB, we inserted a LoxP-Stop-LoxP unit 5’ to RFP. As expected, RFP positive cells were virtually absent 10 weeks after injecting this AAV-shPTB vector into the midbrain of wild-type mice (Extended Data Fig. 4a, left). In contrast, when the same AAV vector was injected into midbrain of transgenic mice that express the Cre recombinase under the astrocyte-specific GFAP promoter^20^, RFP was readily detectable (Extended Data Fig. 4a, right).

**Fig. 3.**
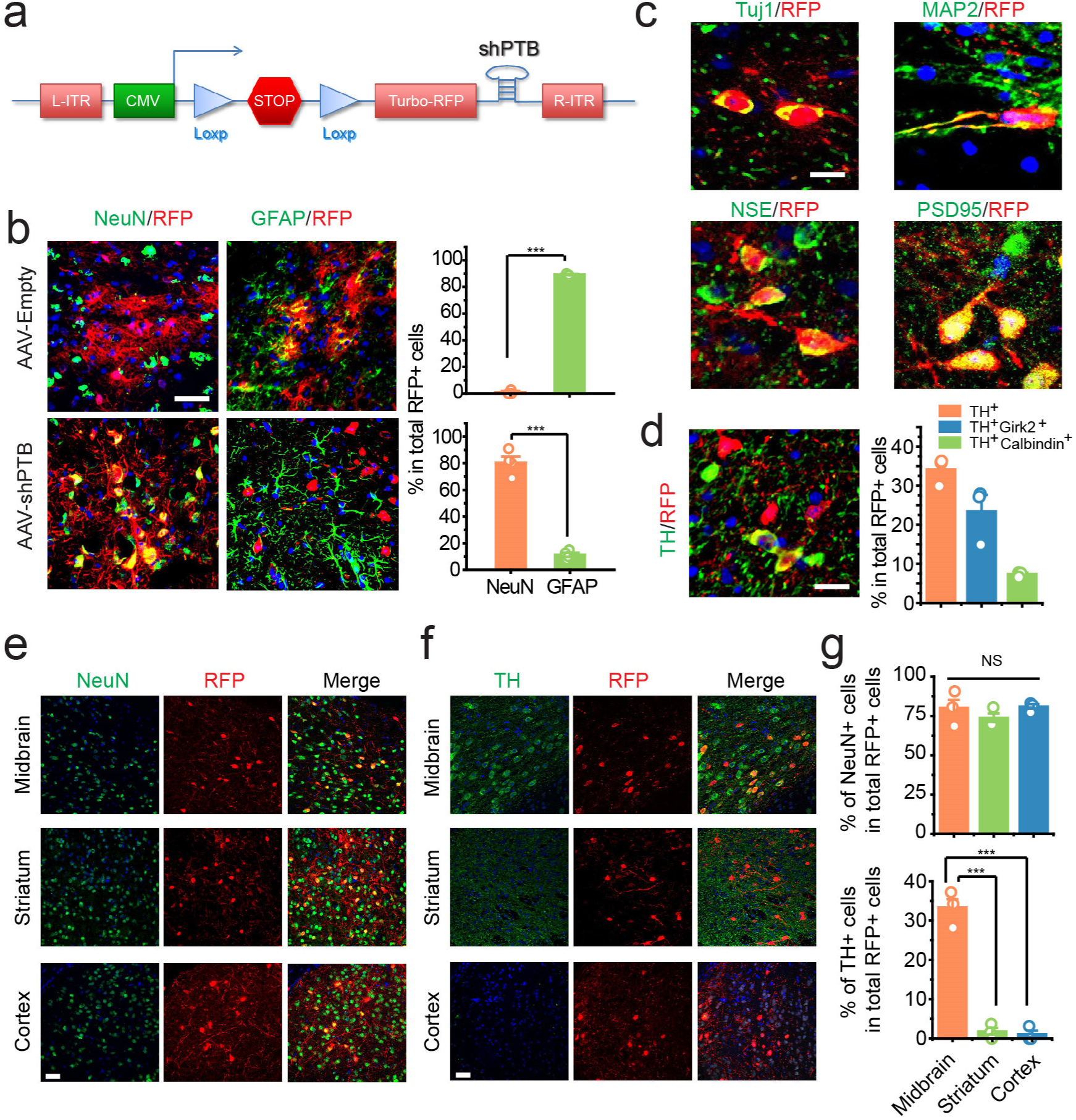
Conversion of astrocytes into mature neurons by PTB down-regulation in different mouse brain regions. **a**, Design of an AAV vector containing two lox-P sites that bracket a transcription stop signal followed by RFP and shRNA against PTB. **b**, AAV-Empty vector-infected astrocytes showed GFAP-positive staining but not NeuN (upper panels), and in contrast, most of astrocytes infected with the AAV-shPTB vector were NeuN positive (lower panels) 10 weeks after transduction. Quantified data were based on 3 mice (AAV-Empty) or 4 mice (AAV-shPTB) as shown on the right. *** p<0.001 based on unpaired Student t-test. Scale bar: 30 μm. **c**, Converted neurons stained for multiple neuronal markers, including Tuj1, MAP2, NSE and PSD95. Scale bar: 10 μm. **d**, A significant population of converted neurons in the midbrain expressed TH. Right panel shows the quantification for subgroups of converted dopaminergic neurons. Quantified data from 3 mice are shown on the right. Scale bar: 20um. Note that the quantitative data are intended to present the actual levels, not for emphasizing statistical differences between different groups. **e** and **f**, AAV-shPTB-induced mature neurons (NeuN) and dopaminergic neurons (TH) in different brain regions as indicated. Scale bar: 30um. **g**, Quantification of the staining results in **e** and **f**. Data were from 3 mice. In panel **b** and **g**: *** p<0.001 based on ANOVA with post-hoc Tukey test.

Next, we chose to inject the AAV-shPTB vector into one side of the substantia nigra of GFAP-Cre mice between P30 and P40, a developmental stage when astrocytes are known to have already lost their neurosphere-generating potential in the midbrain^21^. As a negative control, we performed similar injections with a vector encoding only RFP (AAV-Empty). In the AAV-Empty injected group, as expected, most RFP^+^ cells were GFAP^+^, but NeuN^-^, indicating that none of the transduced astrocytes were converted into a neuron (Fig. 3b, top). In contrast, by 3 weeks post-injection of AAV-shPTB, while most transduced cells remained GFAP^+^ and had typical astrocytic morphology, ∼20% of RFP^+^ cells had activated the expression of the mature neuron marker NeuN. The percentage of these RFP^+^/NeuN^+^ cells more than tripled by 5 weeks (Extended Data Fig. 4b, c), and by 10 weeks, >80% RFP^+^ cells were also NeuN^+^, which no longer expressed detectable GFAP (Fig. 3b, bottom, quantified on the right). We found no evidence that NG2 cells were converted to neurons (Extended Data Fig. 4d). These data indicate that initial RFP^+^ midbrain astrocytes were gradually converted into neurons over a 10-week period of PTB reduction.

### Evidence for regional specificity of neuronal conversion in the brain

Ten weeks after the AAV-shPTB delivery, most converted neurons expressed multiple neuronal markers, including Tuj1, MAP2, NSE and PSD-95 (Fig. 3c). Importantly, staining for PSD-95, a membrane-associated guanylate kinase present in the postsynaptic membrane, showed a typical punctate pattern in RFP^+^/NeuN^+^ cells. Remarkably, in contrast to *in vitro* astrocyte-to-neuron conversion (see Fig. 2c), a substantial portion (∼35%) of RFP^+^ neurons expressed the dopaminergic neuron marker TH (Fig. 3d, left), whereas glutamatergic and GABAergic neurons were rarer (Extended Data Fig. 4e). A significant population of converted cells (∼22%) also expressed Girk2, a marker of A9 dopaminergic neurons, while a minor population (∼7%) was positively stained for calbindin-D28k, a marker of A10 dopaminergic neurons (Extended Data Fig. 4f, g and quantified in Fig. 3d, right). These data implied a regional specificity in *trans*-differentiation of astrocytes into different neuronal subtypes. To further test this possibility, we compared *in vitro* converted astrocytes from cortex versus midbrain, observing that midbrain-derived astrocytes were converted at a much higher (∼10%) efficiency into TH^+^ neurons, compared to only ∼2% conversion of cortex-derived astrocytes as determined both by immunostaining and immunoblotting (Extended Data Fig. 5a-c). These data are consistent with a recent RNA-seq analysis showing that astrocytes from different brain regions exhibited different gene expression programs, suggesting the presence of regionally distinct astrocytes^22^.

We next determined the regional specificity in astrocyte conversion *in vivo*. While astrocytes from three brain regions (midbrain, cortex, and striatum) were converted to NeuN^+^ neurons with a similar high efficiency (∼80%), only astrocytes from midbrain showed ∼35% efficiency in *trans*-differentiation into TH^+^ neurons *in vivo* (Fig. 3e-g). The near absence of astrocyte-derived TH^+^ neurons in the striatum was particularly striking, as this is the region innervated by the axons of nigral dopaminergic neurons. Moreover, the higher percentage of astrocyte-derived TH^+^ neurons converted in midbrain (∼35%) compared to that *in vitro* (∼10%) offers strong evidence that local environmental cues may further enhance converted neurons to develop into specific subtypes in different brain regions. These findings are in line with previous reports that astrocytes from midbrain, but not other brain regions, promote differentiation of neuronal stem cells to dopaminergic neurons^23,24^.

### Integration of midbrain astrocyte-derived neurons into the nigrostriatal pathway

Electrophysiological measurements of astrocyte-converted neurons in the midbrain were performed on brain slices 5∼6 weeks after transduction with AAV-shPTB. RFP^+^ neurons were injected with the fluorescent dye Neurobiotin 488 to mark cells recorded by patch clamp and to assess whether they were dopaminergic based on TH staining (Fig. 4a, b). Typical voltage-dependent currents of Na^+^ and K^+^ channels were detected (Fig. 4c). The neuron also exhibited repetitive action potentials (Fig. 4d) and spontaneous postsynaptic currents (Fig. 4e). These data are evidence that astrocyte-converted TH+ neurons are functional and suggest their integration into the existing neural circuits.

**Fig. 4.**
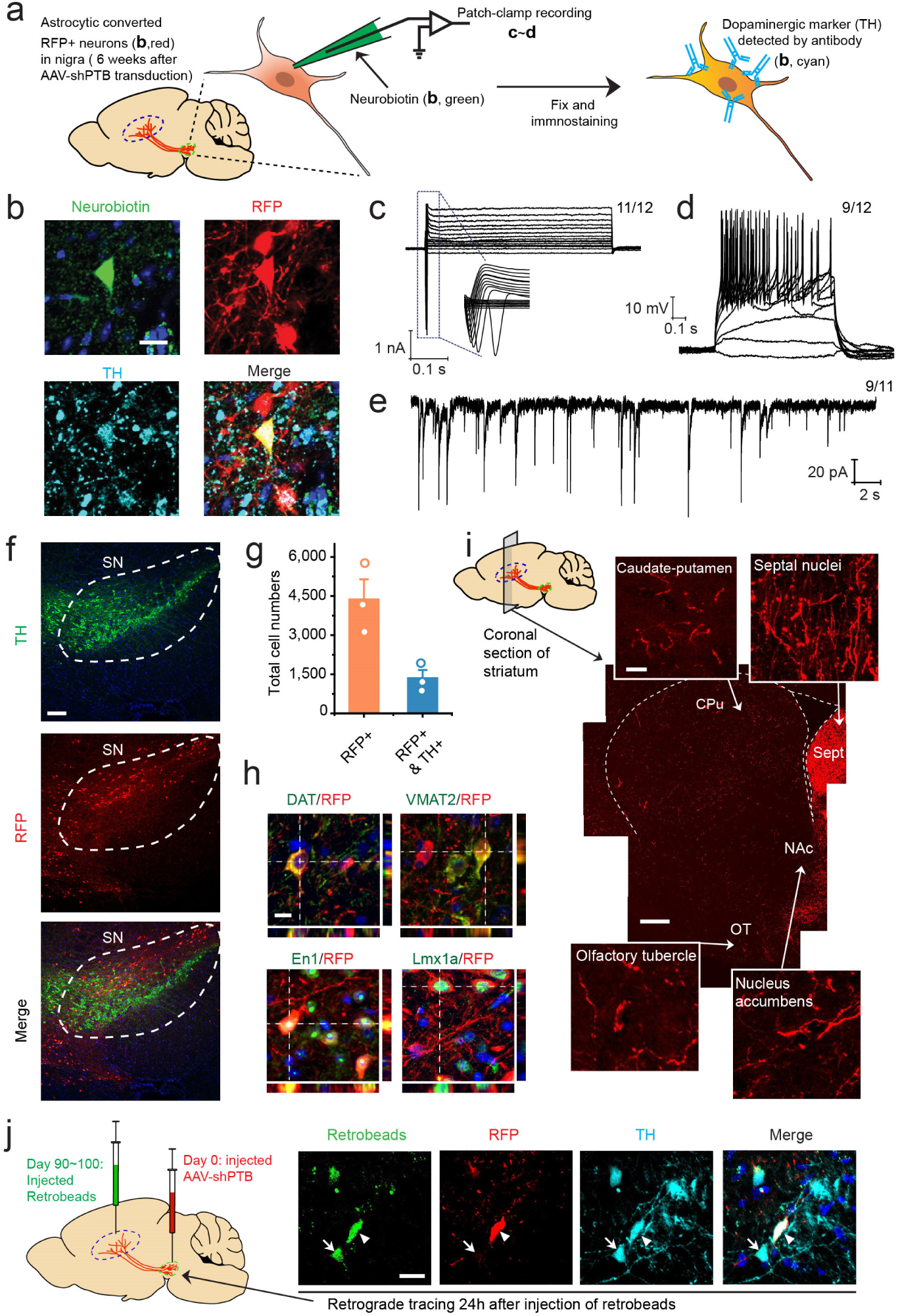
Evidence for integration of midbrain astrocyte-derived neurons into the nigrostriatal pathway. **a**, Schematic depiction of patch recording of converted neurons in the midbrain. **b**, The patched cell was labeled by Neurobiotin 488 filled in the recording pipette. Immunostaining after patch recording demonstrated that the traced cell was TH-positive. Scale bar: 20 μm. **c**,**d**,**e**, Converted neurons on brain slices showed large currents of voltage-dependent sodium and potassium channels (**c**), repetitive action potentials (**d**), and spontaneous post-synaptic currents (**e**). The numbers of cells that showed the recorded activity versus the total number of cells examined are indicated on top right in each panel. **f**, Low magnification view of substantia nigra 10 weeks after injection with AAV-shPTB, stained for TH in comparison with RFP-labeled cells. Scale bar: 100 μm. Note a large number of endogenous TH-positive dopaminergic neurons. RFP-positive cells were detected within substantia nigra (approximated by dashed line) as well as in surrounding regions. **g**, Quantification of total RFP-positive cells in comparison with converted dopaminergic neurons double positive for both RFP and TH. Data were from 3 mice. Note that the quantitative data are intended to present the actual levels, not for emphasizing statistical differences between different groups. **h**, Staining of converted cells for additional markers of dopaminergic neurons, including DAT (dopamine transporter), VMAT2 (vesicular monoamine transporter 2), En1 (engrailed homeobox 1) and Lmx1a (LIM homeobox transcription factor 1 alpha). RFP and TH double positive cell body is highlighted by orthogonal views of z-stack images, attached on right and bottom of the main image in each panel. Scale bar: 10um. **i**, Low magnification view of the striatum innervated by RFP-positive projections. Scale bar: 300um. Inserted panels show the amplified views of RFP-positive projections in different regions. CPu: caudate-putamen; NAc: nucleus accumbens; Sept: septal nuclei; OT: olfactory tubercle. Scale bar: 15 μm. **j**, Labeling of a TH/RFP-double positive cell in substantia nigra with retrograde beads. Arrowhead indicates a beads-labeled converted cell and arrow points to a beads-labeled endogenous dopaminergic neuron (TH-positive but RFP-negative). Scale bar: 20 μm.

Counting of astrocyte-converted neurons revealed ∼4000 total RFP^+^ cells within substantia nigra, of which ∼1300 were TH^+^ neurons (Fig. 4f, g). Subtype specificities of these reprogrammed neurons were further confirmed by immunostaining on brain slices for the dopamine transporter (DAT), the vesicular monoamine transporter 2 (VMAT2), as well as specific markers for midbrain dopaminergic neurons, such as engrailed homeobox 1 (En1) and LIM homeobox transcription factor 1 alpha (Lmx1a) (Fig. 4h). RFP^+^ cell bodies were present only in the substantia nigra, but RFP^+^ fibers were detected in the caudate-putamen (CPu) as well as other target fields, including nucleus accumbens (NAc), septal nuclei (Sept) and the olfactory tubercle (Fig. 4i), as observed in earlier studies in which the mouse midbrain was grafted with neuronal stem cells^25^. A fraction of these fibers were also TH^+^ (Extended Data Fig. 6a). Quantification of fiber density (see Methods) revealed that RFP^+^/TH^+^ processes were mainly distributed in the caudate-putamen and nucleus accumbens regions of the striatum, despite the presence of ∼3-fold more RFP^+^ fibers in septal nuclei (Extended Data Fig. 6b). These data suggest the influence of environmental cue(s) in the direction of innervation by astrocyte-converted neurons.

To further test if axons of newly converted neurons extended from the substantia nigra to the striatum, 3 months after AAV-shPTB delivery we injected green fluorescent retrobeads into the caudate-putamen of mice to allow for axonal uptake and retrograde labeling of the corresponding cell bodies (Fig. 4j). One day after injection, green beads were detected in both endogenous TH^+^/RFP^-^ cells and converted TH^+^/RFP^+^ cells within the substantia nigra (arrows and arrowheads in Fig. 4j, right panel). Thus, both anterograde and retrograde tracing data strongly support the integration of AAV-shPTB reprogrammed dopaminergic neurons into the nigrostriatal pathway.

### Replenishing lost dopaminergic neurons in the nigrostriatal pathway

The numbers of astrocytes converted to dopaminergic neurons and the relatively robust growth of their axons to striatum suggested that PTB down-regulation converted neurons might be able to partially reconstitute an injured nigrostriatal pathway. To explore this possibility, we induced degeneration of dopaminergic neurons through unilateral injection of 6-hydroxydopamine (6-OHDA), a dopamine analog toxic to dopaminergic neurons, into the medial forebrain bundle^26^ (Fig. 5a). As expected, one month after 6-OHDA injection, unilateral loss of TH^+^ cell bodies in the midbrain was observed along with striatal denervation (Fig. 5b). Accompanying the loss of dopaminergic neurons in the lesioned nigra was a dramatically increased population of GFAP^+^ astrocytes (Fig. 5c), indicative of an expected reactive astrocytic response^27^.

**Fig. 5.**
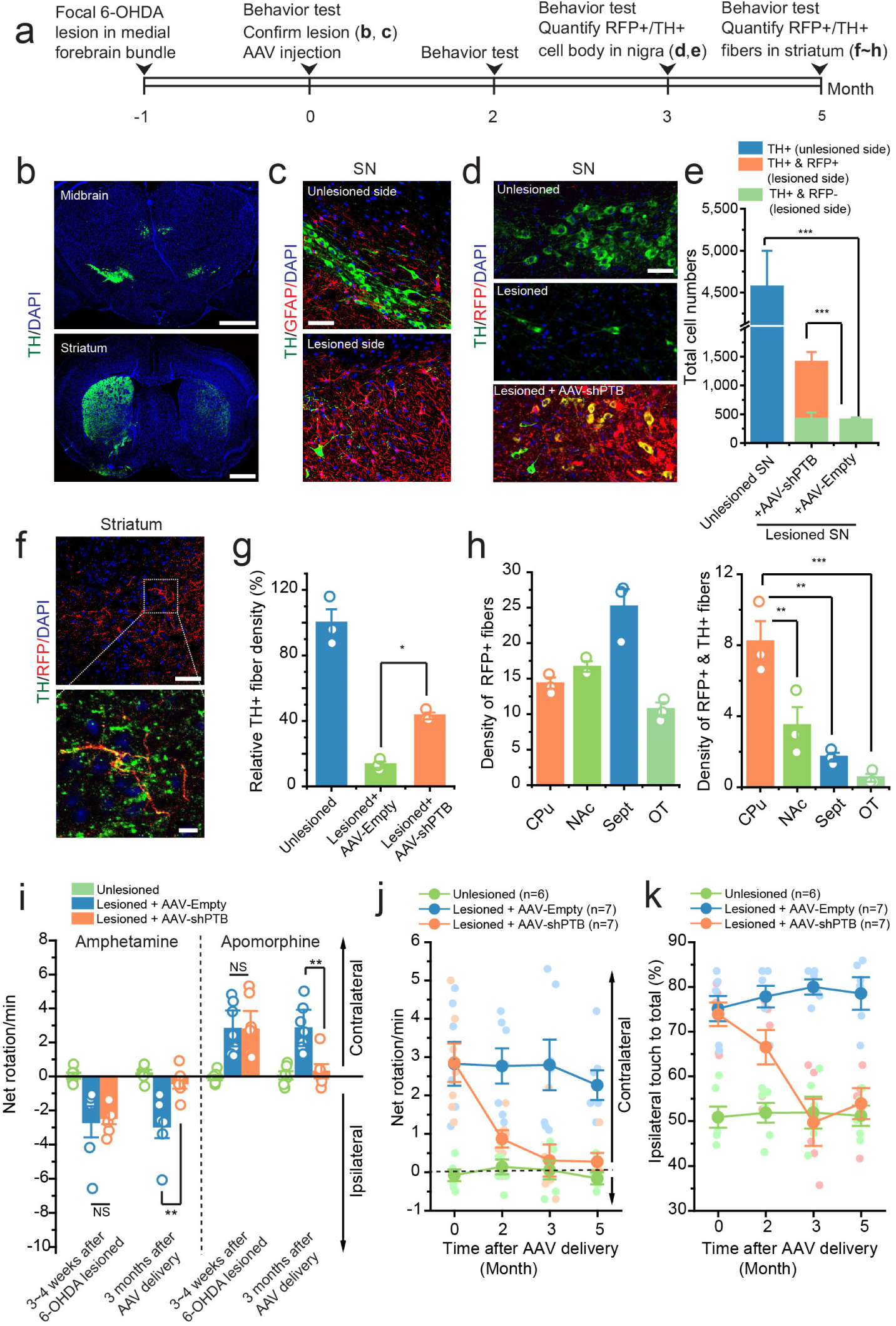
Evidence for reconstruction of the nigrostriatal pathway in a chemically-induced mouse PD model and rescue of PD phenotype. **a**, Schematic depiction of the experimental schedule for 6-OHDA-induced lesion in the substantia nigra (SN), reprogramming with empty vector or AAV-shPTB, behavioral test, and quantification of RPF^+^/TH^+^ cell bodies in SN and fibers in striatum. **b**, Induction of unilateral loss of TH-positive cell bodies in 6-OHDA-induced lesion in the midbrain (top, Scale bar: 500 μm) and TH-stained fiber bundles in the striatum (bottom, Scale bar: 500 μm). **c**, The population of GFAP-positive astrocytes dramatically increased in lesioned nigra. Scale bar: 50 μm. **d**, Comparison between control (top) and 6-OHDA lesioned nigra (middle), showing an increase in converted dopaminergic neurons (yellow) with AAV-shPTB (bottom). Scale bar: 40 μm. **e**, Quantification of dopaminergic, niagral neurons in the unlesioned side (blue), the remaining population of endogenous dopaminergic neurons in the lesioned side (green), and converted RFP/TH double-positive dopaminergic neurons in the lesioned side (orange). Data were from two sets of 3 mice transduced with AAV-shPTB or AAV-Empty vector. **f**, Regenerated RFP/TH-double positive fibers in the striatum. Scale bar: 50 μm (top); 10 μm (bottom). **g**, Quantification for optical densities of total TH-positive fibers in the striatum of unlesioned mice and lesioned mice transduced with AAV-shPTB, showing the restoration of TH-positive dopaminergic neurons in lesioned brain. Data are based on analysis of 3 mice in each group. **h**, Quantification for densities of total RFP-positive fibers and RFP/TH-double positive fibers in 6-OHDA lesioned brain after transduction with AAV-shPTB. Data are based on images from 3 mice. Note that the quantitative data on the left are intended to present the actual levels, not for emphasizing statistical differences between different groups. **i**, Restoration of behavior in mock-treated, 6-OHDA-treated, AAV-shPTB reprogrammed mice. Rotation was induced by amphetamine (left, based on the data from 6 mice) or apomorphine (right, based on the data from 7 mice). **j** and **k**, Time course analysis of behavioral restoration in mock-treated, 6-OHDA-treated, AAV-shPTB reprogrammed mice. Rotation was induced by apomorphine (**j**) and the percentage of ipsilateral touches (**k**) in unilateral lesioned mice was recorded. n=mice analyzed in each group. Statistical results are represented as mean+/- SEM; significant differences are indicated by the p-values based on ANOVA with post-hoc Tukey test. In panel **e, g, h**, and **i**: * p<0.05; **p<0.01; ***p<0.001.

One month after unilateral lesion with 6-OHDA, we injected AAV-Empty or AAV-shPTB into the midbrain. Examination of the lesioned substantia nigra 10∼12 weeks after injection of AAV-shPTB, but not AAV-Empty, showed an increased number of TH^+^ neurons with all of the increase attributable to new, RFP^+^ neurons (Fig. 5d and Extended Data Fig. 7a-d). Neuronal counting revealed that the initial ∼4500 TH^+^ neuronal cell bodies seen in the unlesioned substantia nigra were reduced >90% (to ∼400) in the lesioned side. Importantly, AAV-PTB administration induced ∼1000 new RFP^+^/TH^+^ neurons (Fig. 5e), thereby restoring TH^+^ neurons to ∼1/3 of the initial number. RFP^+^ fibers in the striatum and along the nigrostriatal pathway were detected, a fraction of which was also TH^+^ (Fig. 5f and Extended Data Fig. 8a-f). Quantitative analysis of fiber density indicated that 6-OHDA reduced TH^+^ fibers to ∼15% of the initial level and AAV-PTB restored TH^+^ fibers to ∼40% of wild-type levels in the unlesioned side (Fig. 5g). As in unlesioned brain treated with AAV-shPTB (Extended Data Fig. 6), quantification of RFP^+^ and RFP^+^/TH^+^ fibers in different striatum regions again revealed that the caudate-putamen (CPu) region contained the highest proportion of RFP^+^/TH^+^ fibers (Fig. 5h). Although a similar, partially reconstituted nigrostriatal pathway has been achieved with transplantation of stem cell-derived dopaminergic neurons in mouse brain^25,28,29^, our data now demonstrate that, without any additional treatment to specify neuronal subtypes, AAV-shPTB is able to induce new neurons from endogenous midbrain astrocytes to partially replenish lost dopaminergic neurons.

### Reversal of Parkinson disease phenotype by direct reprogramming in midbrain

To determine the ability to restore motor function to 6-OHDA lesioned mice following PTB suppression to convert astrocytes into new neurons, we performed three commonly used behavior tests, two based on drug-induced rotation and the third on spontaneous motor activities^30-33^. Both contralateral rotation induced by apomorphine and ipsilateral rotation triggered by amphetamine were markedly increased following 6-OHDA induced lesion. Remarkably, both of these phenotypes were restored to nearly wild-type levels within 3 months after AAV-shPTB treatment. No corrections were recorded in AAV-Empty transduced mice (Fig. 5i). Progressive recovery over 2-3 months of apomorphine-induced rotations was recorded in the same set of mice (Fig. 5j).

To examine spontaneous motor activity, we scored limb use bias^30-32^. Unlesioned mice used both limbs with relatively equal frequency, while unilaterally lesioned mice showed preferential ipsilateral touches, indicating disabled contralateral forelimb function. In lesioned mice transduced with AAV-shPTB, there was a time-dependent improvement in contralateral forelimb use, reaching wild-type levels of performance by 3 months post treatment, while AAV-Empty transduced mice failed to show any improvement (Fig. 5k).

To test if the reprogrammed neurons were directly responsible for the restoration of motor function, we employed a chemogenetic approach (Fig. 6a). We replaced RFP in our AAV-shPTB vector with a gene encoding a previously engineered inhibitory muscarinic receptor variant hM4Di, which no longer responds to acetylcholine and remains inactive unless bound by clozapine-N-oxide (CNO)^34^. Correspondingly, neurons converted from astrocytes will incorporate this receptor into their plasma membrane, and CNO activation of hM4Di results in G_i_ signaling that opens potassium channels^35^, thereby decreasing firing of action potentials by any neuron expressing it. CNO is metabolized within 2 to 3 days after injection, thereby restoring the function of converted hM4Di-expressing neurons^34^.

**Fig. 6.**
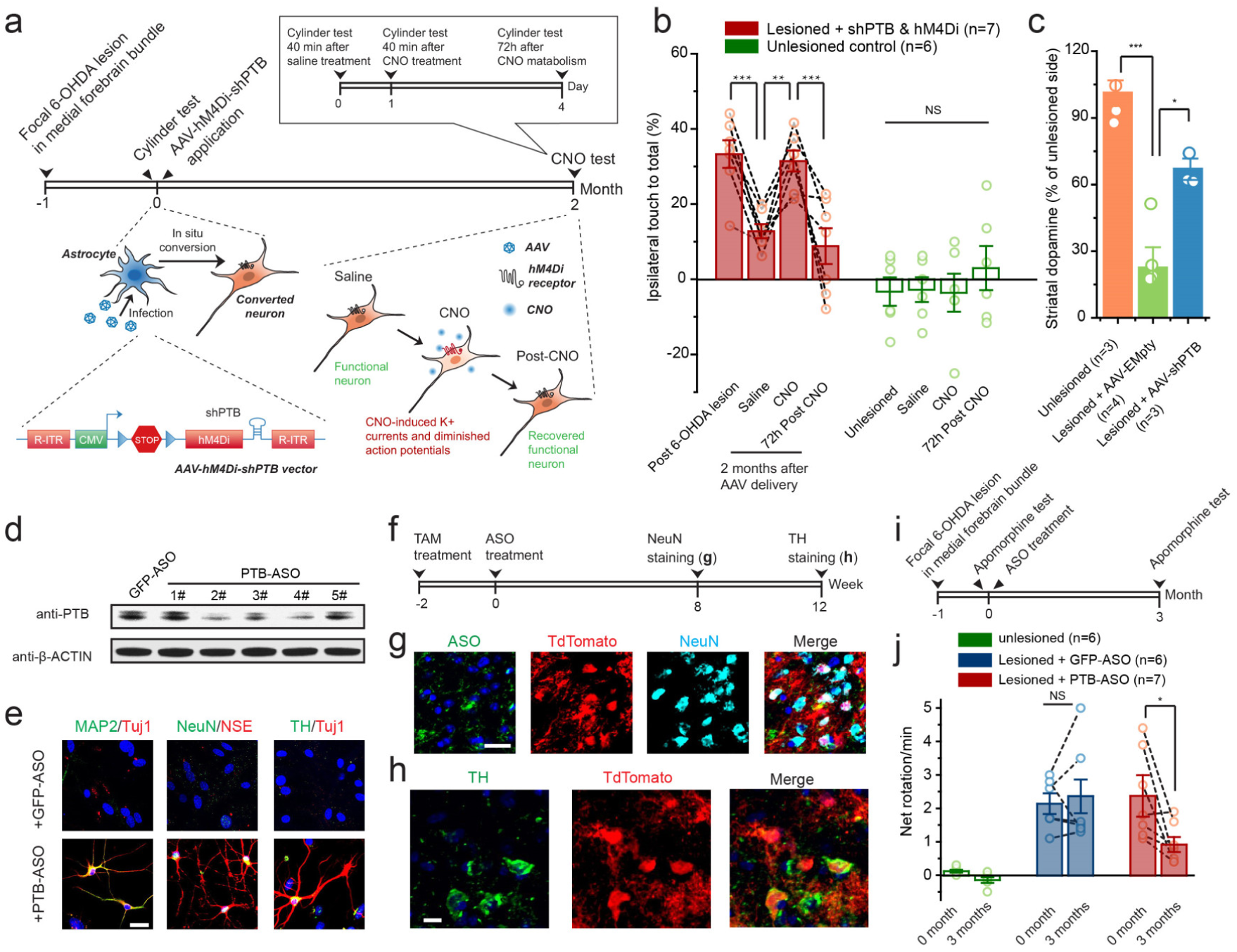
Reversal of Parkinson disease phenotype by direct reprogramming in midbrain. **a**, Schematic depiction of the chemogenetic approach to demonstrate converted neurons are directly responsible for the phenotypic recovery, emphasizing the rapid effect (40 min) of injected CNO to prevent action potentials of reprogrammed neurons and functional restoration after metabolism of the drug in 3 days. **b**, The results of the cylinder test, showing preferential ipsilateral touches in lesioned mice before and after injecting AAV-hM4Di-shPTB as well as treatment with CNO and 3 days after drug withdrawal. Unlesioned mice served as control. n=mice analyzed in each group. Dashed lines show the same mice at different treatment conditions. **c**, Quantification of striatal dopamine levels, showing significant restoration of striatal dopamine after reprogramming with AAV-shPTB in ipsilateral nigra. n=mice analyzed in each group. **d**, Screening for efficient antisense oligonucleotides that target PTB. PTB levels were examined by immunoblotting in mouse astrocytes treated with different ASOs. ASO 4# was chosen for the rest of experiments. **e**, PTB-ASO induced neurons from mouse cortical astrocytes in vitro, which were stained positively for Tuj1 and MAP2 (left), NSE and NeuN (middle), as well as the dopaminergic neuron marker TH (right). Note that as in shPTB-treated cells, a small percentage of TH-positive cells was also detected in PTB-ASO treated astrocytes. Mouse astrocytes treated with GFP-ASO showed no positive staining of any of the neuronal markers under same culture conditions. Scale bar: 20 μm. **f**, Schematic depiction of Tamoxifen (TAM)-induced tdTomato expression and ASO-induced neuronal conversion from tdTomato-labeled astrocytes in substantia nigra. **g**, A fraction of tdTomato-labeled cells became NeuN^+^ 8 weeks after injection of PTB-ASO. Scale bar: 20 μm. **h**, A fraction of PTB-ASO converted tdTomato-positive neurons were TH^+^ in 12 weeks. Scale bar: 10 μm. **i**, Schematic depiction of the experimental schedule for 6-OHDA-induced lesion followed by reprogramming with ASO and behavioral test. **j**, PTB-ASO, but not GFP-ASO, partially rescued the rotation behavior induced by apomorphine in 6-OHDA lesioned mice. Dashed lines show the same mice before and after ASO-mediated reprogramming. In panel **b, c**, and **j**: Statistical results are represented as mean+/- SEM; significant differences are indicated by the p-values based on ANOVA with post-hoc Tukey test (b and c) or paired t-test. *p<0.05; **p<0.01; ***p<0.001.

As expected, motor performance of 6-OHDA lesioned mice was restored (measured with the limb bias assay) within 2 months of transduction of astrocytes with the AAV-hM4Di-shPTB virus (Fig. 6b). Within 40 min of intraperitoneal injection of CNO but not saline, the lesioned phenotype re-appeared (Fig. 6b). Injection of CNO into unlesioned mice had no effect. Moreover, the CNO-provoked motor phenotype disappeared within 3 days, correlating with expected metabolism of the chemical (Fig. 6b). The rapid return and disappearance of the PD-like phenotype in CNO-treated mice thus provide strong evidence that electrical signaling by astrocyte-derived neurons is responsible for phenotypic recovery and suggests limited contribution of trophic factor(s) secreted by converted neurons that led to the repair of damaged endogenous neurons.

### Restoration of striatal dopamine in reprogrammed brain

Given the very rapid CNO-induced phenotypic return and then its loss within 3 days, we next asked whether AAV-shPTB-induced neurons restored dopamine in the striatum. To this end, extracts were prepared from the striatum and dopamine levels were quantified by HPLC analysis. Known amounts of spike-in dopamine were used to identify its elution position and show a linear correlation between signal intensity and the amount of spiked dopamine (Extended Data Fig. 9a-b). Levels of dopamine in the striatum from both lesioned and unlesioned sides of mouse brain were then measured with or without AAV-shPTB mediated astrocyte conversion, each based on 3 independent experiments. 6-OHDA lesioning reduced dopamine to ∼25% of the normal level (Fig. 6c and Extended Data Fig. 9c-f). Remarkably, after AAV-shPTB treatment, dopamine was elevated 250% compared to its level in the lesioned striatum, reaching 65% of the unlesioned level (Fig. 6c). This ∼40% net gain is similar to the 30 to 35% recovery of RFP+/TH+ neuronal cell bodies in the nigra and processes in the striatum, strongly suggesting that AAV-shPTB reprogrammed neurons are responsible for the observed phenotypic recovery.

### ASO-induced neuronal conversion and rescue of chemically induced PD

The PTB-regulatory loop can be self re-enforcing once triggered by initial PTB knockdown (because PTB itself is a target of miR-124 - see Fig. 1a): In response to PTB reduction, miR-124 becomes more efficient in targeting REST (due to the ability of PTB to directly compete with the miRNA targeting site in the 3’UTR of *REST*), resulting in reduced REST, which drives further de-repression of miR-124; as REST is an established repressor of this miRNA, elevation in miR-124 further suppresses PTB expression^11,16^. Recognition of this regulatory circuit enabled a strategy for transient suppression of PTB through antisense oligonucleotide (ASO)-mediated degradation by intranuclear RNase H of PTB mRNAs bound to a complementary ASO. This approach has been shown to mediate target mRNA degradation in rodent and non-human primate nervous systems after injection into the cerebral spinal fluid^36-38^.

We therefore synthesized and screened 21-nucleotide anti-PTB ASOs that contained a phosphorothioate backbone (to increase metabolic stability and aid with delivery) with a 3’ fluorescein (to allow tracing of the injected ASO). We also synthesized a control ASO targeting GFP. Three PTB-ASOs, but not the GFP-ASO, were identified to be effective in reducing PTB expression upon transfection of isolated mouse astrocytes (Fig. 6d). Five weeks after introduction of the PTB-ASO with the highest targeting efficiency, mouse astrocytes cultured in standard neuronal differentiation medium induced synthesis of a series of neuronal markers, including Tuj1, MAP2, NSE and NeuN, while astrocytes transfected with the control GFP-ASO did not (Fig. 6e). A small fraction of converted neurons were dopaminergic (a proportion similar to that produced by AAV-shPTB), as indicated by positive TH staining (Fig. 6e).

Whether transient ASO-mediated suppression of PTB could induce astrocyte conversion into neurons in the midbrain was tested in mice carrying a Cre-recombinase encoding transgene which upon induction with tamoxifen will be selectively expressed from the astrocyte-specific GFAP promoter and a tdTomato encoding gene (integrated at the Rosa26 locus) whose expression will be permanently activated by the action of the Cre recombinase. We systemically induced Cre at postnatal day 35 (P35) (Extended Data Fig. 10a) and 2 weeks later stereotactically injected ASOs unilaterally into the substantia nigra. Without injected ASO, tdTomato-labeled cells were all NeuN^-^ (Extended Data Fig. 10b). In contrast, by 2∼3 months post PTB-ASO administration, a portion of tdTomato-labeled cells became NeuN^+^ (Fig. 6f, g) and TH^+^ (Fig. 6h). Unilateral 6-OHDA-lesioning produced apomorphine-induced rotations, and by three months post 6-OHDA-lesioning, the PTB-ASO, but not control GFP-ASO, produced significant phenotypic rescue (Fig. 6i, j). These findings illustrate a highly clinically feasible approach to the treatment of PD.

## DISCUSSION

In this report, we demonstrate that astrocytes can be reprogrammed *in vivo* by reducing the level of a single RNA binding protein. Our approach is fundamentally distinct from TF-based reprogramming, which requires ectopic expression of an exogenous gene(s). Indeed, it simplifies the approach by creating a method that employs a single step to transiently suppress a single endogenous gene. Our strategy exploits the existence of a regulatory loop in which transient down-regulation of PTB leads to a self-sustaining off switch^16^. Our approach also exploits the discovery that the genetic underpinnings of a neuronal differentiation program are present, but latent, in mouse and human astrocytes and that once fully induced by suppressing PTB produces mature neurons. Importantly, our findings define a clinically feasible approach to generate neurons from local astrocytes in mammalian brain using a single dose of an anti-PTB ASO. This approach now enables efforts to define its effectiveness in reconstituting vulnerable neuronal circuits in neurodegenerative disorders.

The phenotype(s) of PTB knockdown-induced neurons appears to be a function of the context in which they are produced and/or the astrocytes from which they are derived. While much effort has been applied using TF-based approaches to generate specific types of neurons^39-41^, our findings point to a powerful role for the *in vivo* environment to define the results of reprogramming. Consistent with recent RNA-seq profiling^22^, our data reveal that astrocytes of different brain regions are intrinsically distinct, with midbrain-derived astrocytes particularly prone to conversion into dopaminergic neurons. Furthermore, we find that the local environment, likely defined in part by the presence of regionally distinct astrocytes, further enhances development of specific neuronal subtypes and the maintenance of their axons in normal targets of innervation. These findings are consistent with the documented effects of midbrain-derived astrocytes on inducing differentiation of co-cultured neuronal stem cells into dopaminergic neurons^23,24^.

Our data indicate that PTB reduction in the mammalian brain converts astrocytes to dopaminergic neurons capable of restoring dopamine biogenesis, accompanied by reversal of behavioral deficits in a chemically-induced PD model. Although it has been a subject of debate whether this model recapitulates essential features of PD pathogenesis^42^, it does result in a critical endpoint, the loss of substantia nigra neurons and depletion of striatal dopamine. Significantly, the chemogenetic experiments to transiently inactivate electrical signaling of the newly induced neurons demonstrate that those neurons are directly responsible for the benefits at the behavioral levels due to at least partial reconstitution of the nigrostriatal pathway, as judged by the presence within the substantia nigra of newly converted neurons and in striatum of the processes from these neurons. However, we cannot rule out the contribution of any tropic factor(s) secreted by astrocyte-derived neurons or potential fusion between new neurons and the remaining endogenous neurons that led to the overall recovery of the disease phenotype.

Our approach thus meets all five generally accepted milestones set for *in vivo* reprogramming^8^. That said, much work is now required to establish efficacy of astrocyte-to-neuron conversion, to determine how fully the nigrostriatal pathway can be reconstituted by the newly generated neurons, and to explore its potential use as a therapy for disorders of the human nervous system. Important objectives include the ability to specifically target regions harboring vulnerable neurons, inducing an effective but not excessive number of neurons exhibiting desired physiological phenotypes and capable of releasing desired neurotransmitter(s), optimizing synaptic engagement of the processes of converted neurons with appropriate inputs and target neurons, and ensuring the reconstitution of functional properties and behavioral rescue. Importantly, each of these objectives can now be addressed to develop this promising therapeutic strategy.

## Acknowledgement

The authors are grateful to members of the Fu lab for cooperation, reagent sharing, and insightful discussion during the course of this investigation. We thank Dr. Alysson Muotri of UCSD for a gift of human ES cell-derived neural progenitors. D.W.C. receives salary and research support from the Ludwig Institute for Cancer Research and is a Nomis Foundation Distinguished Scientist. This work was supported by NIH grants (GM049369 and GM052872) to X.D.F.

## Supplementary Materials

Materials and Methods

Extended Data Figure 1 to 10

## METHODS

### Vectors and virus production

To build the lentiviral vector to express shPTB in mouse astrocytes, the target sequence (5’-GGGTGAAGATCCTGTTCAATA-3’) was shuttled to the pLKO.1-Hygromycin vector (Addgene, #24150). For human astrocytes, a similar vector containing target sequence (5’-GCGTGAAGATCCTGTTCAATA-3’) was used. Viral particles were packaged in Lenti-X 293T cells (Clontech) with two package plasmids: pCMV-VSV-G (Addgene, #8454) and pCMV-dR8.2 dvpr (Addgene, #8455). Viral particles were concentrated by ultracentrifugation on a Beckman XL-90 centrifuge with SW-28 rotor.

To construct AAV vectors, the same target sequence against mouse PTB was first inserted into the pTRIPZ-RFP vector between EcoR I and Xho I sites. The segment containing RFP and shRNA was next sub-cloned to replace CaMP3.0 in AAV-CMV-LOX-STOP-LOX-mG-CaMP3.0 vector (Addgene, #50022) by using Asc I. To construct a control vector, a similar segment containing only RFP was cloned into the AAV-CMV-LOX-STOP-LOX-mG-CaMP3.0 vector. The resulting vectors were referred to as AAV-shPTB or AAV-Empty. The AAV-hM4Di-shPTB vector was constructed by replacing RFP in AAV-shPTB with the cDNA of hM4Di, which was sub-cloned from pAAV-CBA-DIO-hM4Di-mCherry vector (Addgene, #81008).

Viral particles of AAV2 were packaged in transfected 293T cells with other two plasmids: pAAV-RC and pAAV-Helper (Agilent Genomics). After harvest, viral particles were purified with heparin column (GE HEALTHCARE BIOSCIENCES) and then concentrated with Ultra-4 centrifugal filter units (Amicon, 100,000 molecular weight cutoff). Titers of viral particles were determined by qPCR to be >1x 10^12^ particles/ml.

### Synthesis of antisense oligonucleotides

Antisense oligonucleotides were synthesized by Integrated DNA Technologies. The sequence of the target region in mouse PTB for ASO synthesis is 5’-GGGTGAAGATCCTGTTCAATA-3’, and the target sequence in Turbo GFP is 5’-CAACAAGATGAAGAGCACCAA-3’. The backbones of all ASOs contain phosphorothioate modifications. Fluorescein (FAM) was attached to 3’ end of those ASOs for fluorescence detection.

### Western blot and RT-PCR

For analysis by western blotting, cells were lysed in 1xSDS loading buffer, and after quantification, bromophenol blue was added to a final concentration of 0.1%. 25∼30ug of total proteins were resolved in 10% Nupage Bis-Tris gel and probed with following antibodies: Rabbit anti-PTBP1 (kindly provided by Douglas Black, 1:3000), Mouse anti-PTBP2 (Santa Cruz, sc-376316, 1:1000), Mouse anti-beta actin (Sigma, A2228, 1:10000),Rabbit anti-Tuj1 (Covance, MRB-435P, 1:10000),Rabbit anti-Brn2 (Cell Signaling, 12137, 1:1000), Chicken anti-TH (Aves lab, TYH, 1:1000) and Rabbit anti-VMAT2 (Proteintech, 20873-1-AP, 1:500).

For RT-qPCR analysis, RNA was extract with Trizol (Life Technology) and 10ug/ml of Glycogen was used to enhance precipitation of small RNAs. Total RNA was first treated with DNase I (Promega) followed by reverse transcription with miScript II RT Kit (QIAGEN, 218160). RT-qPCR was performed using the miScript SYBR Green PCR Kit (QIAGEN, 218073) on a step-one plus PCR machine (Applied Biosystems). The primers used were U6-F: 5’-ACGCAAATTCGTGAAGCGTT-3’; miR-124-F: 5’-TAAGGCACGCGGTGAATGCC-3’; and miR-9-F: 5’-GCGCTCTTTGGTTATCTAGCTGTATG-3’.

### Cell culture and *trans*-differentiation *in vitro*

Mouse astrocytes were isolated from postnatal (P4∼P5) pups. The cortical tissue was dissected from whole brain and incubated with Trypsin before plating onto dishes coated with Poly-D-lysine (Sigma). Isolated astrocytes were cultured in DMEM (GIBCO) plus 10% fetal bovine serum (FBS) and penicillin/streptomycin (GIBCO). Dishes were carefully shaken daily to eliminate non-astrocytic cells. After reaching ∼90% confluency, astrocytes were disassociated with Accutase (Innovative Cell Technologies) followed by centrifugation for 3 min at 800 rpm, and then cultured in medium containing DMEM/F12 (GIBCO), 10% FBS (GIBCO), penicillin/streptomycin (GIBCO), B27 (GIBCO), 10 ng/ml epidermal growth factor (EGF, PeproTech), and 10 ng/ml fibroblast growth factor 2 (FGF2, PeproTech).

To induce *trans*-differentiation *in vitro*, mouse astrocytes were re-suspended with astrocyte culture medium containing the lentivirus that targets mouse PTB, and then plated on Matrigel Matrix (Corning)–coated coverslips (12 mm). After 24 hrs, cells were selected with hygromycin B (100ug/ml, Invitrogen) in fresh astrocyte culture medium for 72 hrs. The medium was next switched to the N3/basal medium (1:1 mix of DMEM/F12 and Neurobasal, 25 μg/ml insulin, 50 μg/ml transferring, 30 nM sodium selenite, 20 nM progesterone, 100 nM putrescine,) supplemented with 0.4% B27, 2% FBS, a cocktail of 3 small molecules (1 μM ChIR99021, 10 μM SB431542 and 1mM Db-cAMP), and neurotrophic factors (BDNF, GDNF, NT3 and CNTF, all in 10ng/ml). The medium was half-changed every the other day. To measure synaptic currents, converted cells after 5∼6 weeks were added with fresh GFP-labeled rat astrocytes, and after further 2 to 3 weeks of co-culture, patch-clamp recordings were performed.

Human astrocytes were purchased from a commercial source (ScienCell). Cells were grown in Astrocyte Medium (ScienCell) and sub-cultured until reaching ∼80% confluency. For *trans*-differentiation *in vitro*, cultured human astrocytes were first disassociated with Trypsin; re-suspended in Astrocyte Medium containing the lentivirus that targets human PTB; and plated on Matrigel Matrix–coated coverslips. After 24 hrs, cells were selected with hygromycin B (100ug/ml, Invitrogen) for 72 hrs. The medium was switched to the N3/basal medium supplemented with 0.4% B27, 2% FBS and neurotrophic factors (BDNF, GDNF, NT3 and CNTF, all in 10ng/ml). To measure synaptic currents, converted cells after 3 weeks were added with fresh GFP-labeled rat astrocytes, and after further 2 to 3 weeks of co-culture, patch-clamp recordings were performed.

### Immunocytochemistry

Cultured cells grown on glass slides were fixed with 4% Paraformaldehyde (Affymetrix) for 15 min at room temperature followed by permeabilization with 0.1% Triton X-100 in PBS for 15 min on ice. After washing twice with PBS, cells were blocked in PBS containing 3% BSA for 1 hr at room temperature. The fixed cells were incubated with primary antibodies overnight at 4°C in PBS containing 3% BSA. After washing twice with PBS, the cells were incubated with secondary antibodies conjugated to Alexa Fluor 488, Alexa 546, Alexa 594 or Alexa 647 (1:500, Molecular Probes) for 1 hr. 300 nM DAPI in PBS was applied to the cells for 20 min at room temperature to label nuclei. After additional washing three times with PBS, Fluoromount-G mounting media was applied onto the glass slides, and images were examined and recorded under Olympus FluoView FV1000.

For staining brain sections, mice were sacrificed with CO2 and immediately perfused, first with 15∼20mL saline (0.9% NaCl) and then with 15 mL 4% paraformaldehyde (PFA) in PBS to fix tissues. Whole brains were extracted and fixed in 4% PFA overnight at 4°C, and then cut to 14∼18um sections by a cryostat (Leica). Before staining, brain sections were incubated with sodium citrate buffer (10 mM Sodium citrate, 0.05% Tween 20, pH 6.0) for 15 min at 95°C for antigen retrieval. The slides were next treated with 5% normal donkey serum and 0.3% Triton X-100 in PBS for 1 hr at room temperature. The rest of steps were performed as on cultured cells.

The following primary antibodies were used: Rabbit anti-Tuj1 (Covance, MRB-435P, 1:1,000), Mouse anti-Tuj1 (Covance, MMS-435P, 1:1,000), Mouse anti-MAP2 (Milipore, MAB3418, 1:1000), Mouse anti-NeuN (Milipore, MAB377, 1:200), Chicken anti-NSE (Aves lab, NSE, 1:1000), Rabbit anti-VGlut1 (Synaptic Systems, 135-303, 1:200), Rabbit anti-GAD67 (Cell Signaling, 63080, 1:200), Rabbit anti-GAD67 (GeneScript, A01440, 1:100), Mouse anti-VGlut1 (BioLegend, MMS-5245-100, 1:200), Chicken anti-TH (Aves lab, TYH, 1:1000), Rabbit anti-PSD95 (Cell Signaling, 3450, 1:200), Rabbit anti-DAT (Bioss, bs-1714R, 1:100), Goat anti-VMAT2 (Everest biotech, EB06558, 1:100), Rabbit anti-VMAT2 (Proteintech, 20873-1-AP, 1:100), Rabbit anti-DAT (Proteintech, 22524-1-AP, 1:100), Rabbit anti-En1 (Abgent, AP7278a, 1:100), Rabbit anti-Lmx1a (ProSci, 7087, 1:100), Rabbit anti-GFAP (Cell Signaling, 12389, 1:200), Chicken anti-GFAP (Aves lab, GFAP, 1:100), Rabbit anti-ALDH1L1 (EnCor Biotechnology, RPCA-ALDH1L1, 1:2000), Mouse anti-OLIG2 (Santa Cruz, sc-293163, 1:100), Chicken anti-CD11b (Aves lab, MAC, 1:1000), Mouse anti-NG2 (Santa Cruz, sc-53389, 1:100), Mouse anti-Nestin (Cell Signaling, 4760, 1:200), Mouse anti-NANOG (Santa Cruz, sc-293121, 1:100), Mouse anti-Fibronectin (DSHB, 1H9, 1:500), Rabbit anti-GAD65 (Cell Signaling, 5843, 1:50), Rabbit anti-VGlut2 (Bioss, bs-9686R, 1:100), Rabbit anti Girk2 (Proteintech, 21647-1-AP, 1:100), Rabbit anti Calbindin D28K (Proteintech, 14479-1-AP, 1:100) and Mouse anti-RFP (ThermoFisher, MA5-15257, 1:200).

### Quantification of neuronal cell body and fiber density

Coronal sections across the midbrain were sampled at intervals of 120∼140 μm for immunostaining of TH and RFP. The total numbers (Nt) of cell types of interest were calculated using the formula of Nt = Ns*(St/Ss) in which Ns is the number of neurons counted, St is the total number of sections in the brain region, and Ss is the number of sections sampled, as previously described^43,44^.

RFP-positive and RFP/TH-double positive fibers were quantified using a previously published method ^25^. Three coronal sections (A/P +1.3, +1.0 and +0.70) were selected from each brain for analysis ^43^. For each selected section, three randomly chosen areas were captured from one section of z-stack images at intervals of 2 µm using a 60x oil-immersion objective. A sphere (diameter: 14 µm) was then generated as a probe to measure fiber density within the whole z-stack. Each fiber crossing the surface of sphere was given one score. The optical density of striatal TH fibers was determined from same sections. The digitalized image of sampled section was captured with a 10x objective and analyzed by Image-J 1.47v (Wayne Rasband, Bethesda, MD)^45,46^.

### Electrophysiology

Patch clamp recordings were performed with Axopatch-1D amplifiers or Axopatch 200B amplifier (Axon Instruments) connecting to a Digidata1440A interface (Axon Instruments). Data were acquired with pClamp 10.0 or Igor 4.04 software and analyzed with MatLab v2009b. For converted neurons from mouse astrocytes *in vitro*, small molecules were removed from medium 1 week before patch clamp recording. Both cultured mouse and human cells were first incubated with oxygenated (95% O2/5% CO2) artificial cerebrospinal fluid (150 mM NaCl, 5 mM KCl, 1 mM CaCl2, 2 mM MgCl2, 10 mM glucose, 10 mM HEPES, pH 7.4) at 37°C for 30 min and whole-cell patch clamp was performed on selected cells.

For recording activities of converted neurons *in vivo*, cortical slices (300 µm) were prepared 5∼6 weeks after injections of AAV vectors. The slices were cut with a vibratome in oxygenized (95% O2/5% CO2) dissection buffer (110.0 mM choline chloride, 25.0 mM NaHCO3, 1.25 mM NaH2PO4, 2.5 mM KCl, 0.5 mM CaCl2, 7.0 mM MgCl2, 25.0 mM glucose, 11.6 mM ascorbic acid, 3.1 mM pyruvic acid) at 4°C followed by incubation in oxygenated ACSF (124 mM NaCl, 3 mM KCl, 1.2 mM NaH2PO4, 26 mM NaHCO3, 2.4 mM CaCl2, 1.3 mM MgSO4, 10 mM dextrose and 5 mM HEPES; pH 7.4) at room temperature for 1 hr before experiments.

Patch pipettes (5–8 MΩ) solution contained 150 mM KCl, 5 mM NaCl, 1 mM MgCl2, 2 mM ethylene glycol tetra acetic acid (EGTA)-Na, and 10 mM Hepes pH 7.2. For voltage-clamp experiments, the membrane potential was typically held at −75 mV. The following concentrations of channel blockers were used: PiTX: 50uM; NBQX: 20uM; APV: 50uM. All of these blockers were bath-applied following dilution into the external solution from concentrated stock solutions. All experiments were performed at room temperature.

### Construction of mouse models

The GFAP-Cre transgenic mouse (B6.Cg-Tg(Gfap-cre)77.6Mvs/2J) was used in AAV-shPTB induced *in vivo* reprogramming experiments. For testing the effect of ASOs *in vivo*, the GFAP-CreER™ mouse (B6.Cg-Tg(GFAP-cre/ERT2)505Fmv/J) was crossed with the Rosa-tdTomato mouse (B6.Cg-Gt(ROSA)26Sortm14(CAG-tdTomato)Hze/J). The offsprings of these double GFAP-CreER™;Rosa-tdTomato transgenic mice aged postnatal 35 days were injected with tamoxifen (dissolved in corn oil at a concentration of 20 mg/ml) via intraperitoneal injection once every 24 hrs for a total of 5 consecutive days. The dose of each injection was 75 mg/kg. Two weeks after tamoxifen application, PTB-ASO or control ASO was injected into the substantia nigra of those mice to investigate ASO-induced *in vivo* reprogramming.

All transgenic mice were purchased from The Jackson Laboratory. All procedures were conducted in accordance with the guide of The University of California San Diego Institutional Animal Care and Use Committee (Protocol# S99116).

### Ipsilateral lesion with 6-OHDA and stereotaxic injections

Adult WT and GFAP-Cre mice at the age of postnatal day 30 were used to perform surgery to induce lesion. Animals were first anaesthetized with a mix of ketamine (80-100mg/kg) and xylazine (8-10mg/kg) and then placed in a stereotaxic mouse frame. Before injecting 6-hydroxydopamine (6-OHDA, Sigma), mice were first treated with a mix of desipramine (25 mg/kg) and pargyline (5mg/kg). 6-OHDA was dissolved in 0.02% ice-cold ascorbate/saline solution at a concentration of 15mg/ml and used within 3 hrs. The toxic solution was injected into the medial forebrain bundle (MFB) at the following coordinates (relative to bregma): anterior–posterior (A/P) = −1.2mm; medio-lateral (M/L) = −1.3mm and dorso-ventral (D/V) = −4.75mm (from the dura). Injections were applied in a 5 ul Hamilton syringe with a 33G needle at the speed of 0.1ul/min for 3 min before slowly removing the needle. Cleaning and suturing of the wound were performed after lesion.

AAV vectors or ASOs were injected into the substantia nigra ∼30 days after 6-OHDA induced lesion. 4ul of AAV vectors or 2ul of ASOs (1ug/ul) were injected into lesioned nigra at the following coordinates A/P = −3.0mm; M/L = −1.2mm and D/V = −4.5 mm. Injections were made using same syringe and needle, at a rate of 0.5ul/min for 3 min before slowly removing the needle.

### Retrograde tracing

For retrograde tracing of nigrostriatal pathway, GFAP-Cre mice with or without 6-OHDA induced lesion were first injected with AAV-shPTB vectors. 3 months after the AAV delivery, green Retrobeads IX (Lumafluor, Naples, FL) were unilaterally injected at two sites into the striatum on the same side of AAV injection, using following two coordinates: A/P = + 0.5mm, M/L = + 2.0mm, D/V= + 3.0mm and A/P= + 1.2mm, M/L= + 2.0mm, D/V= + 3.0mm. ∼2ul of beads were injected. After 24 hrs, animals were sacrificed and immediately perfused. Their brains were fixed with 4%PFA for sectioning and immunostaining.

### Measurement of striatal dopamine

Dopamine levels in mouse striatum were measured by Reverse-phase High-performance Liquid Chromatography (HPLC). The HPLC analysis was performed using an Agilent 1260 Infinity HPLC system with an Agilent Zorbax SB-C18 semi-prep column (ID 9.4 x 250 mm, 5 μm, 80Å) using a water/methanol gradient containing 0.1% formic acid. Each substance is characterized by retention time and 260 nm absorbance under Variable Wavelength Detector (VWD), as previously described ^47,48^. The striatal samples were directly prepared from brain tissue. Briefly, striatal dissection was carried out immediately after euthanization of the mouse. After homogenized in 200 µL of 0.1M hydrochloric acid with a squisher, the sample was centrifuged (12,000 × g, 10 min, 4°C). The resulting supernatant was filtered by a 0.2um Nanosep MF centrifugal device and then applied to HPLC analysis^48,49^.

### Behavioral testing

All behavioral tests were carried out 21-28 days after 6-OHDA induced lesion or 2, 3, and 5 months after the delivery of AAV vectors or ASOs. For rotation test, apomorphine-induced rotations in mice were recorded after intraperitoneal injection of apomorphine (Sigma, 0.5mg/kg) under a live video system. Mice were injected with apomorphine (0.5mg/kg) on two separate days prior to performing the rotation test (for example, if the test was to be performed on Friday, the mouse would be first injected on Monday and Wednesday), which aimed to prevent a ‘wind-up’ effect that could obscure the final results. Rotation was measured 5 min following the injection for 10 min periods as previously described ^31,33^ and only full-body turns were counted. Rotations induced by D-amphetamine (Sigma, 5mg/kg) were determined in the same system ^32,50^. Data are expressed as net contralateral or ipsilateral turns per min. For cylinder test, mice were individually placed into a glass cylinder (diameter 19 cm, height 25 cm), with mirrors placed behind for a full view of all touches, as described^30,31^. Mice were recorded under a live video system for 5 min. No habituation of the mice to the cylinder was performed before the recording. A frame-by-frame video player (KMPlayer version 4.0.7.1) was used for scoring. Only wall touches independently with the ipsilateral or the contralateral forelimb were counted. Simultaneous wall touches (touched executed with both paws at the same time) were not included in the analysis. Data are expressed as a percentage of ipsilateral touches in total touches. For chemogenetic experiment, cylinder tests were carried out 21-28 days after 6-OHDA induced lesion and 2 months after the delivery of AAV-hM4Di-shPTB. In the later test, animal was firstly injected with saline to record the baseline of recovery. Subsequent recording was performed 40 min after Intraperitoneal injection of CNO (Biomol International, 4 mg/kg) or 72 hrs after metabolism of the drug^35^.

### Data analysis and statistics

The exact numbers (n) of biological replicates or mice are indicated in individual figure legends. Experimental variations in each graph are represented as mean+/- SEM. All measurements were performed on independent samples. Independent t-test, one-way ANOVA and repeat measurement ANOVA were employed for statistical analysis, as indicated by individual figure legends. For multiple comparisons, combining ANOVA, post-hoc Tukey test was applied. Assumptions of normal data distribution and homoscedasticity were adopted in independent t-tests and one-way ANOVA. All statistical tests were two-tailed. For Fig. 1b, Fig. 1c and Fig. 5e, the original data were transformed to their logarithms with base 10 for one-way ANOVA to fulfill the requirement of homoscedasticity.

