## Supplemental Figures 1 to 10 for "Therapeutic Reversal of Chemically Induced Parkinson Disease by Converting Astrocytes into Nigral Neurons"

### EXTENDED DATA FIGURES AND LEGENDS

Extended Data Figure 1

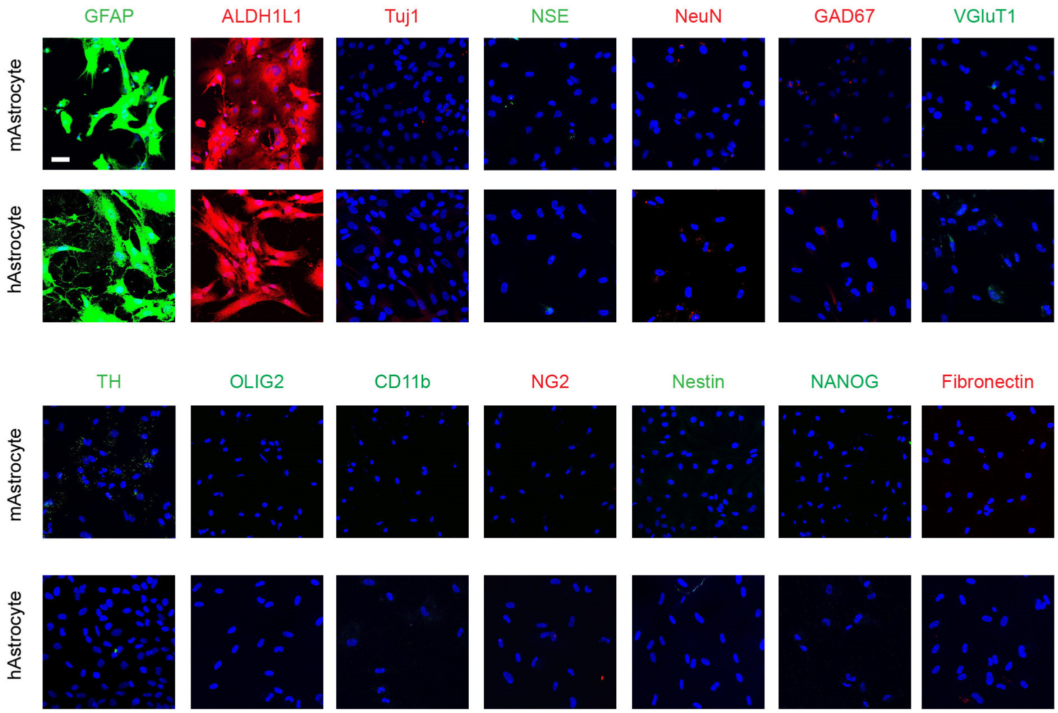


**Extended Data Fig. 1 | Characterization of isolated mouse and human astrocytes**
The majority of mouse and human astrocytes in our culture were immunopositive for astrocyte markers (GFAP and Aldehyde Dehydrogenase 1 Family Member L1, ALDH1L1), without detectable other cell types, as indicated by negative staining of neuronal markers (Tuj1, NSE, NeuN, GAD67, VGluT1, TH), oligodendrocyte marker (Oligodendrocyte Transcription Factor 2, OLIG2), Microglia marker (CD11 Antigen-Like Family Member B, CD11b), NG2 cell marker (Neural/glial antigen 2, NG2), neural progenitor marker (Nestin) pluripotency marker (NANOG) and fibroblast marker (Fibronectin). Scale bar: 30 μm.

Extended Data Figure 2

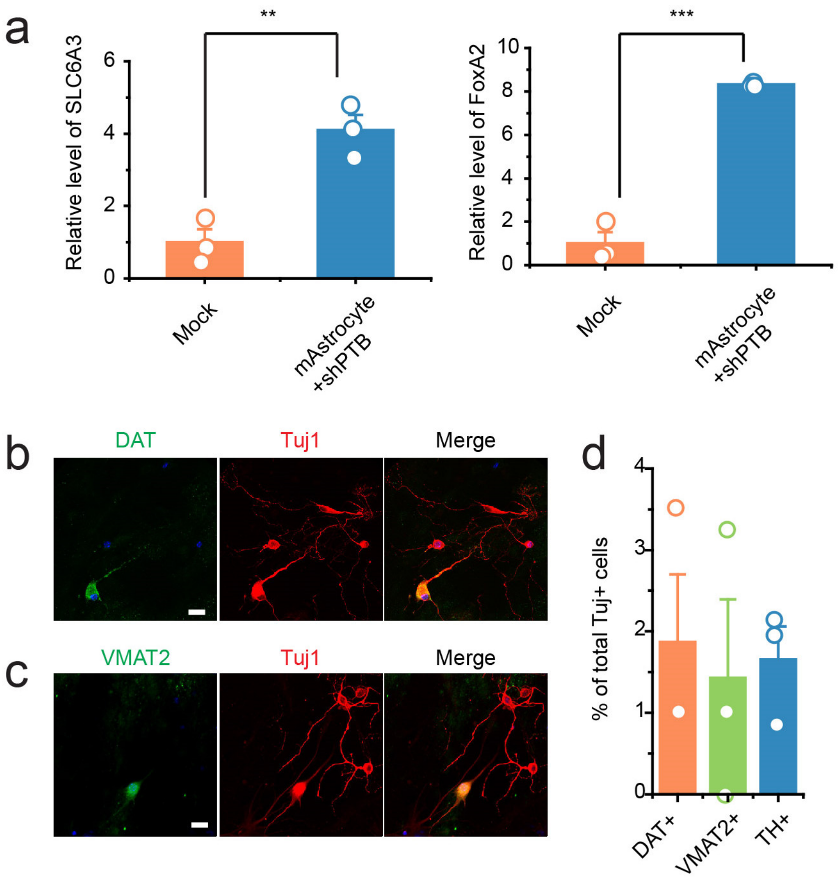


**Extended Data Fig. 2 | Expression of dopaminergic neuron markers in cortical astrocyte-derived neurons in response to PTB knockdown *in vitro.* a,** Expressions levels of SLC6A3 (left) and FoxA2 (right) measured by RT-qPCR in mouse cortical astrocytes before and after conversion to neurons with by shPTB. Statistical significance was determined by Student's t-test based on 3 biological repeats and represented as mean+/- SEM. **p<0.01; ***p<0.001. **b** and **c**, Characterization of induced dopaminergic neurons by immunostaining for DAT (**b**) and VMAT2 (**c**). Scale bar: 20 µm. **d**, Quantification of the percentage of converted neurons that express DAT and VMAT2 in comparison with TH. Data were based on 3 biological replicates in each case.

Extended Data Figure 3

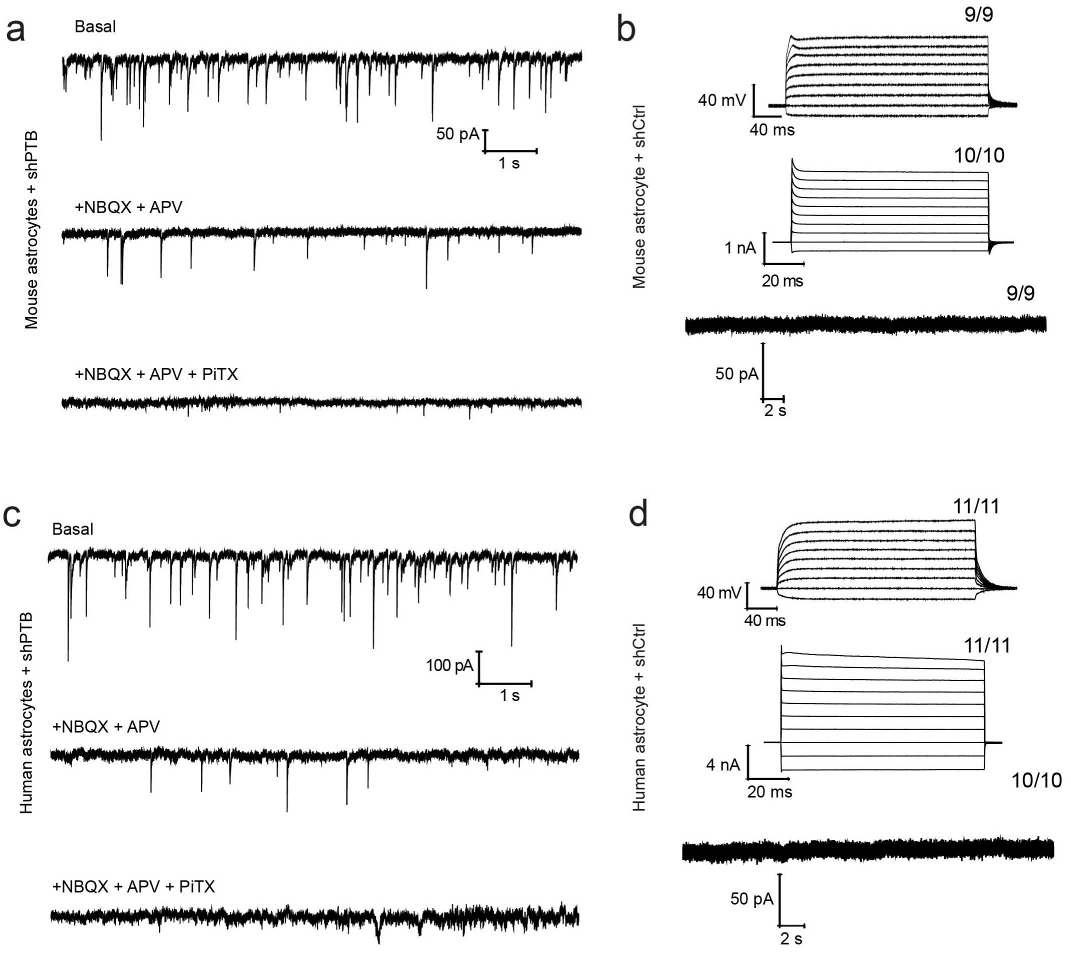


**Extended Data Fig. 3 | Electrophysiological properties of converted neurons and mock-treated astrocytes. a** and **c**, Spontaneous excitatory and inhibitory postsynaptic currents were detected both on mouse (**a**) and human (**c**) astrocytes after PTB knockdown-induced neuronal conversion. The currents could be sequentially blocked by inhibitors against the excitatory (NBQX+APV) and inhibitory (PiTX) receptors. **b** and **d**, Mock mouse (**b**) and human astrocytes (**d**) did not show any neuronal electrophysiological properties, such as action potentials (top), currents of voltage-dependent channels (middle) and postsynaptic events (bottom). The numbers of cells that showed the recorded activity versus the total number of cells examined are indicated on top right in each panel.

Extended Data Figure 4

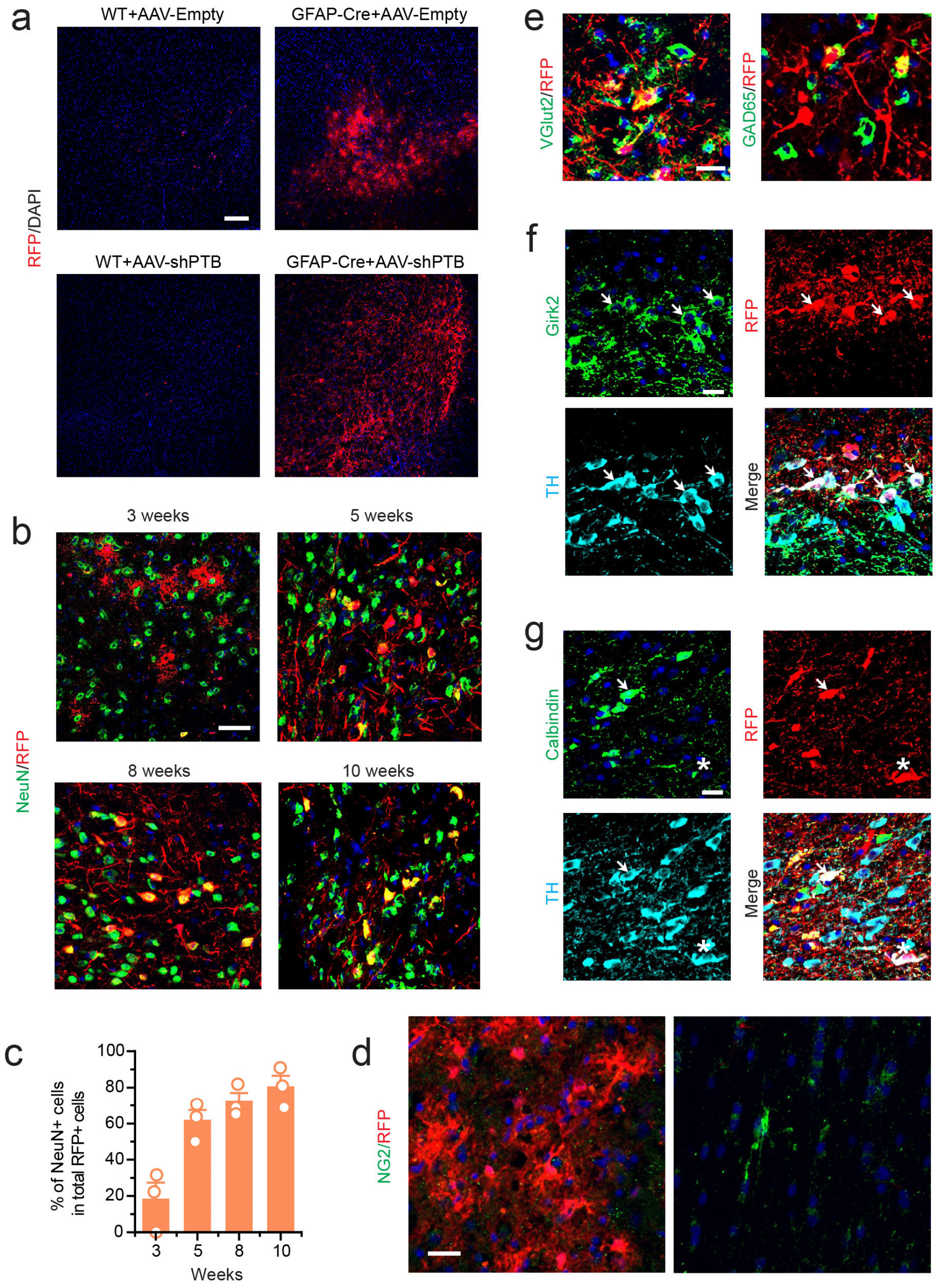


**Extended Data Fig. 4 | AAV-shPTB induced neuronal conversion in the mouse midbrain. a,** Undetectable leakage of the LoxP-Stop-LoxP AAV expression unit in injected wild-type mice. RFP-positive cells were rarely detected in the midbrain of wild-type mice (left) 10 weeks after injection of AAV-Empty or AAV-shPTB virus, in comparison with GFAP-Cre transgenic mice that had received the same viral injection (right). Scale bar: 150 μm. **b**, RFP-positive cells were gradually converted to neurons in the midbrain. The percentage of RFP-labeled NeuN-positive cells was progressively increased from 3 to 10 weeks post-injection of AAV-shPTB. Scale bar: 50 μm. **c,** Quantification of the data in (b), each based on 3 mice. **d,** NG2 cells were rarely detected around RFP-positive cells (left panel) whereas NG2-positive cells were in general not surrounded by RFP-positive cells (right panel) in the same slice of AAV-Empty transduced midbrain. Scale bar: 15 μm. **e,** Immunostaining of glutamatergic neuron marker (VGluT2) and GABAergic neuron marker (GAD65) showed that different subtypes of converted neurons. Scale bar: 20 μm. **f,** Immunostaining of the A9 dopaminergic neurons marker Girk2. Arrows indicate co-localization of Girk2 with RFP and TH stained signals. Scale bar: 20 μm. **g,** Immunostaining of the A10 dopaminergic neurons marker Calbindin. Arrows indicate co-localization of calbindin with RFP and TH signals. Star indicates a converted dopaminergic neuron (RFP/TH-double positive) that stained negatively for Calbindin. Scale bar: 20 μm.

Extended Data Figure 5

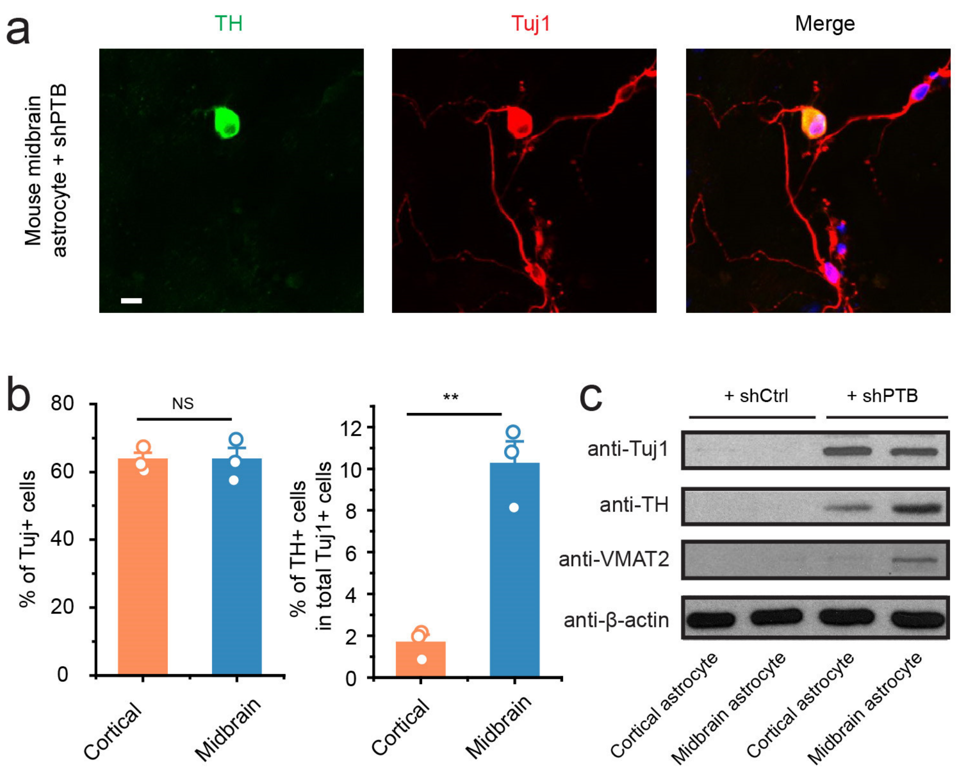


**Extended Data Fig. 5 | The efficiency in converting cortical and midbrain astrocytes to dopaminergic neurons by shPTB *in vitro.* a,** TH staining of shPTB-converted Tuj1-positive neurons from astrocytes derived from midbrain. Scale bar: 10 μm. **b**, Comparison of neuronal conversion efficiencies between astrocytes derived from cortex and midbrain, showing similar high percentage of Tuj1-positive neurons (left), but a significantly higher percentage of dopaminergic neurons converted from midbrain-derived astrocytes relative to that from cortex (right). Statistical significance was determined by Student's t-test based on 3 biological repeats and represented as mean+/-SEM. **p<0.01. **c**, Expression of a pan-neuronal marker (Tuj1) and two specific markers for dopaminergic neurons (TH, VMAT2) in astrocyte-derived neurons from cortex and midbrain, highlighting much higher levels of dopaminergic neuron markers expressed in neurons derived from midbrain astrocytes.

Extended Data Figure 6

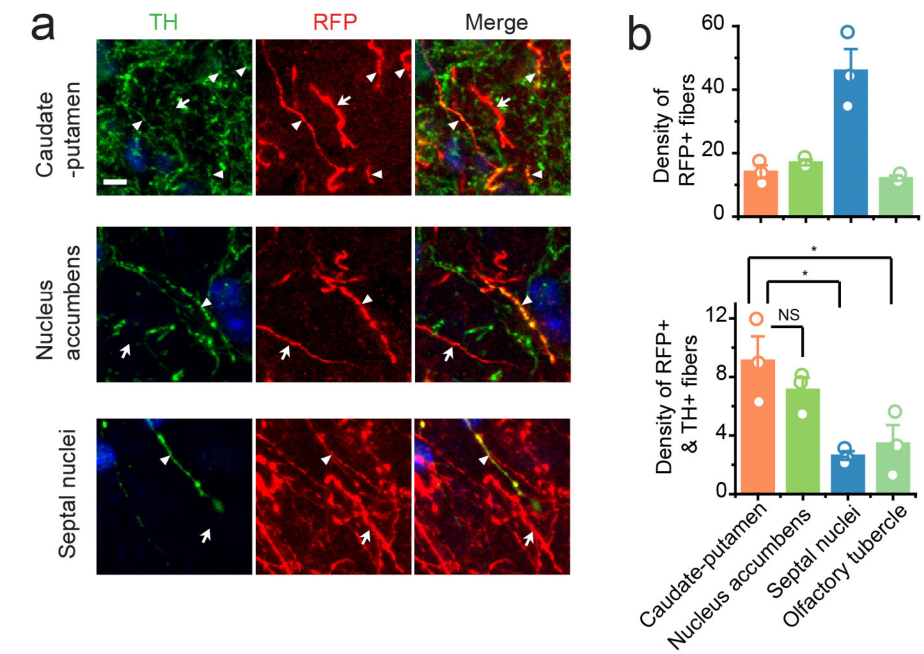


**Extended Data Fig. 6 | shPTB-converted dopaminergic neurons targeted multiple regions. a**, A fraction of RFP-positive fibers showed co-staining with TH (arrowheads), while others are TH-negative (arrows). Scale bar: 5um. **b**, Quantification of densities of total RFP-positive fibers and RFP/TH-double positive fibers after transduction of AAV-shPTB in wild-type mouse brain. Data are based on images from three mice. Statistical results are based on ANOVA with post-hoc Tukey test. * p<0.05.

Extended Data Figure 7

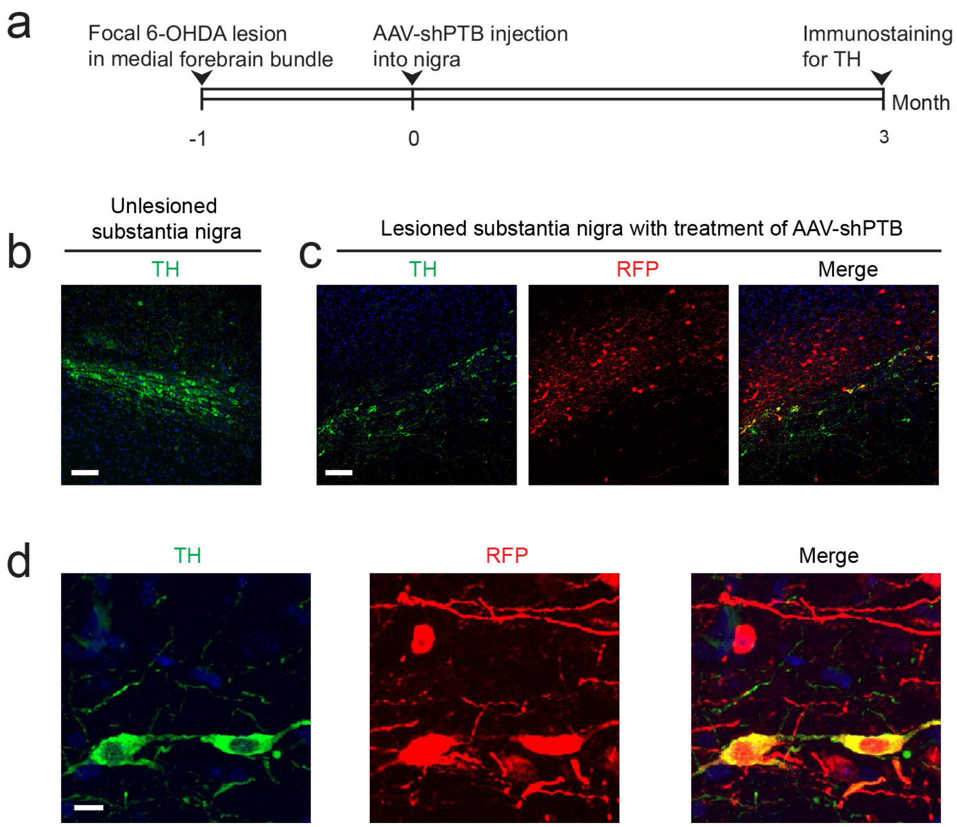


**Extended Data Fig. 7 | shPTB-converted neurons replenish a significant fraction of lost dopaminergic neurons in substantia nigra. a,** Schematic depiction of the experimental schedule for 6-OHDA-induced lesion followed by reprogramming with AAV-PTB. **b,** Low magnification view of unlesioned substantia nigra stained for TH. Scale bar: 80 µm. **c**, The nigra lesioned with 6-OHDA and transduced with AAV-shPTB. Scale bar: 80 µm. The nigra lesioned with 6-OHDA but treated with empty viral vector looked identical between lesioned but untreated nigra (not shown). **d**, An enlarged view of RFP-positive cells also expressing TH in substantia nigra. Scale bar: 10 µm.

Extended Data Figure 8

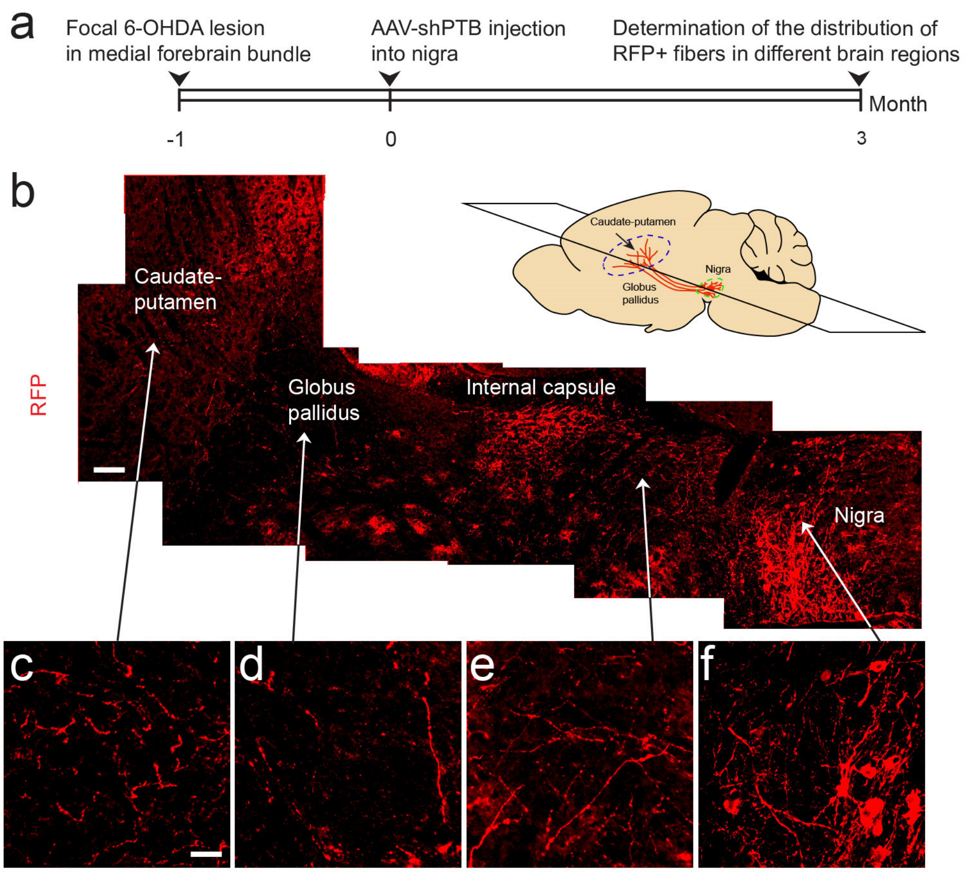


**Extended Data Fig. 8 | Reconstruction of the nigral-striatal pathway by converted dopaminergic neurons. a,** Schematic depiction of the experimental schedule for 6-OHDA-induced lesion and reconstruction of pathway. **b**, Images of RFP-positive projections extended from nigra to striatum. Schematic figure shows the dorso-ventral level of horizontal section. Scale bar: 100 µm. **c** to **f**, Higher magnification views of different brain regions. Scale bar: 25um.

Extended Data Figure 9

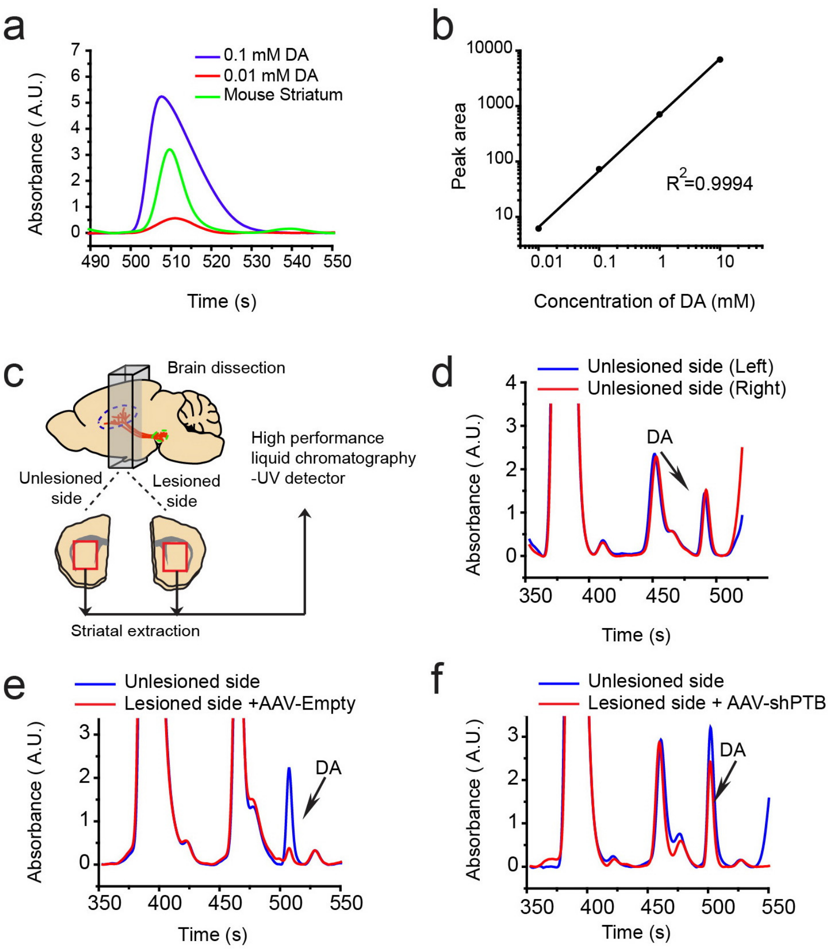


**Extended Data Fig. 9 | Measurement of striatal dopamine by HPLC. a,** Dopamine levels in brain detected by HPLC after two different doses of "spike-in" dopamine" added within the range of dopamine in wild-type brain. **b**, A standard curve generated by "spike-in" dopamine added to different levels. **c**, Schematic depiction of the measurement of striatal dopamine by HPLC. **d**, Comparison of striatal dopamine levels in two sides of the unlesioned mouse brain. **e**, Reduction of striatal dopamine in response to unilateral 6-OHDA lesion. **f**, Significant restoration of striatal dopamine after reprogramming with AAV-shPTB in ipsilateral nigra.

Extended Data Figure 10

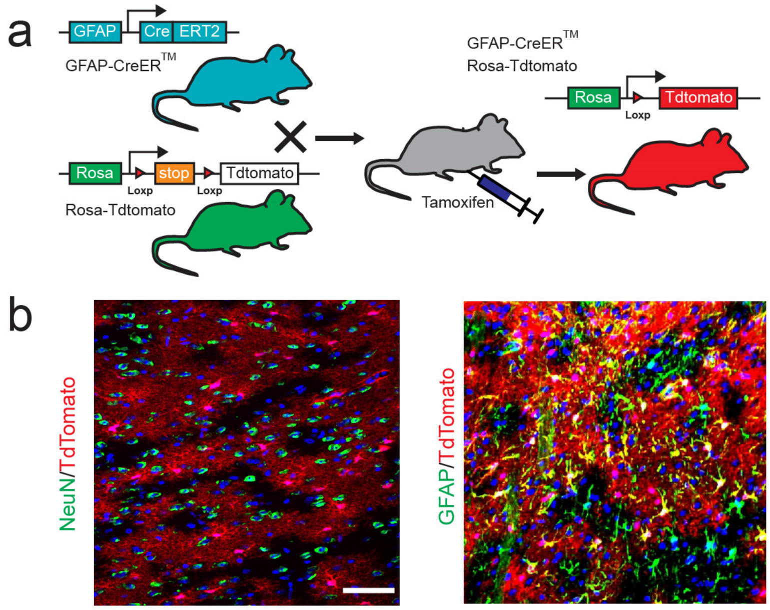


**Extended Data Fig. 10 | PTB-ASO induced neuronal conversion of astrocyte in mouse midbrain. a,** Schematic depiction of transgenic mice used to trace astrocytes *in vivo*. **b**, In the midbrain of the double transgenic GFAP-CreER$^{TM}$; Rosa-tdTomato mouse, 3 weeks after treatment of tamoxifen, none of tdTomato-labeled cells were stained positive for NeuN (left), but most of them were GFAP-positive (right). Scale bar: 50 µm.
